# Mre11-Rad50 enhances spacer acquisition in a haloarchaeal Type I-B CRISPR-Cas system

**DOI:** 10.64898/2026.05.08.723854

**Authors:** Alice K. Cassel, Héloïse Carion, Luciano A. Marraffini

## Abstract

Clustered regularly interspaced short palindromic repeat (CRISPR) loci and their associated (*cas*) genes provide adaptive immunity to bacteria and archaea. CRISPR-Cas systems acquire short DNA fragments from the genomes of infecting plasmids and viruses, which are inserted into the CRISPR locus as a “spacer” sequence in between repeats. Spacers constitute a memory of infection that is used to recognize and attack invading genetic elements in future infections. Despite the evolutionarily divergent genetic backgrounds of bacteria and archaea, the same CRISPR-Cas systems are functional in both of these prokaryotic domains. In bacteria, efficient spacer acquisition requires the DNA repair nucleases RecBCD/AddAB. These nucleases, however, are not present in archaea. Here we investigated the importance of the DNA repair systems in the *Haloferax volcanii* Type I-B CRISPR-Cas response. We found that elimination of the DNA repair nuclease Mre11-Rad50, but not Fen1, substantially reduces spacer acquisition. CRISPR immunity against *H. volcanii* pleomorphic virus 1 (HFPV-1), on the other hand, was not affected by these deletions. Our results describe how CRISPR-Cas systems have adapted to provide anti-viral defense to hosts from different domains of life.

## INTRODUCTION

Bacteria and archaea harbor clustered, regularly interspaced, short palindromic repeat (CRISPR) loci that are flanked by CRISPR-associated genes (*cas*) (1). These loci encode for CRISPR-Cas systems that provide adaptive immunity through the integration of short DNA sequences referred to as “spacers” derived from the genomes of foreign elements such as plasmids (2) and viruses (3) in between the repetitive sequences of the CRISPR array. Spacer acquisition is an exceedingly rare event, occurring in only one of 10^5^-10^6^ cells (4–6) and allows the host organism to recognize the invader in future infections. Spacers acquired from invading agents are fixed into the CRISPR locus due to their defensive properties, but next generation sequencing studies revealed that spacers derived from the host genome can also be incorporated (a process known as self-acquisition), but are most likely lost from the population due to their genotoxicity (7–10). Once integrated into the CRISPR array, spacers are transcribed and processed into short guide RNAs known as CRISPR RNAs (crRNAs) that guide a Cas protein complex to complementary sequences derived from the invader, the “protospacer”, to trigger the CRISPR-Cas immune response (11–13). Depending on the *cas* gene content, these systems can be classified into six types (1). Types I, II, IV and V use the guide RNA to find complementary DNA sequences, a recognition that results in the degradation or the prevention of the replication of the genome of the invader (14). In contrast, during type III and VI CRISPR-Cas immunity, the crRNA recognizes a complementary sequence in an invader’s transcript and triggers a halt in the replication of the host that prevents the spread of the infecting plasmid or virus (15). While some of the earliest descriptions of CRISPR loci were in archaea (16), the bulk of mechanistic CRISPR research has since been done in bacteria.

*Haloferax volcanii* is an established archaeal model organism with a genome composed of a main chromosome and three chromosomal plasmids (pHV1, pHV3, and pHV4) (17), that harbors a Type I-B CRISPR-Cas system (16, 18, 19) with three CRISPR arrays and eight *cas* genes (Fig. 1A). As is the case for all CRISPR-Cas systems, new spacers are inserted into the CRISPR array by the Cas1-Cas2 integrase complex (5, 20–22). Cas4 is also involved in spacer acquisition, but its exact role remains unknown (23). Both repeats and spacers within the CRISPR array are transcribed as a long precursor crRNA (pre-crRNA), which is cleaved into crRNAs by the Cas6b endonuclease (24). Cas6b associates with the Cascade complex (composed of eight Cas7, two Cas5, and one Cas8b subunits) to load the cleaved crRNA guide (11, 24). This complex scans DNA molecules for the presence of a protospacer adjacent motif (PAM) and tests for complementarity between the guide crRNA and the DNA sequence immediately downstream of this motif (25, 26). In the case of base pairing, the Cascade complex recruits Cas3, an helicase/exonuclease that degrades the invader’s genome at the target site (27). Targeting can also influence spacer acquisition through a process known as “primed adaptation”. As opposed to “naïve adaptation” which occurs in the absence of preexisting targets for the spacers of the CRISPR array, primed spacer acquisition is mediated by a spacer with complete or partial homology to the invading genome that is inefficient to provide immunity but leads instead to high rates of spacer acquisition in the vicinity of the target site (4, 10, 28). In *H. volcanii*, primed adaptation was investigated in the presence of low and high concentrations of a self-targeting crRNA produced from a low- or high-copy number plasmid, respectively (29). Spacers were acquired from the entire chromosome and plasmids with hotspots in transposase and rRNA genes and origins of replication, but at high concentrations the most distinct hotspot was located at the self-targeting sites (29). In the absence of the self-targeting spacer, naïve adaptation occurred at very low levels and required over-expression of Cas1, Cas2 and Cas4 for efficient detection, however the overall levels of self-acquisition were very low and no hotspots were determined (29).

**Figure 1.**
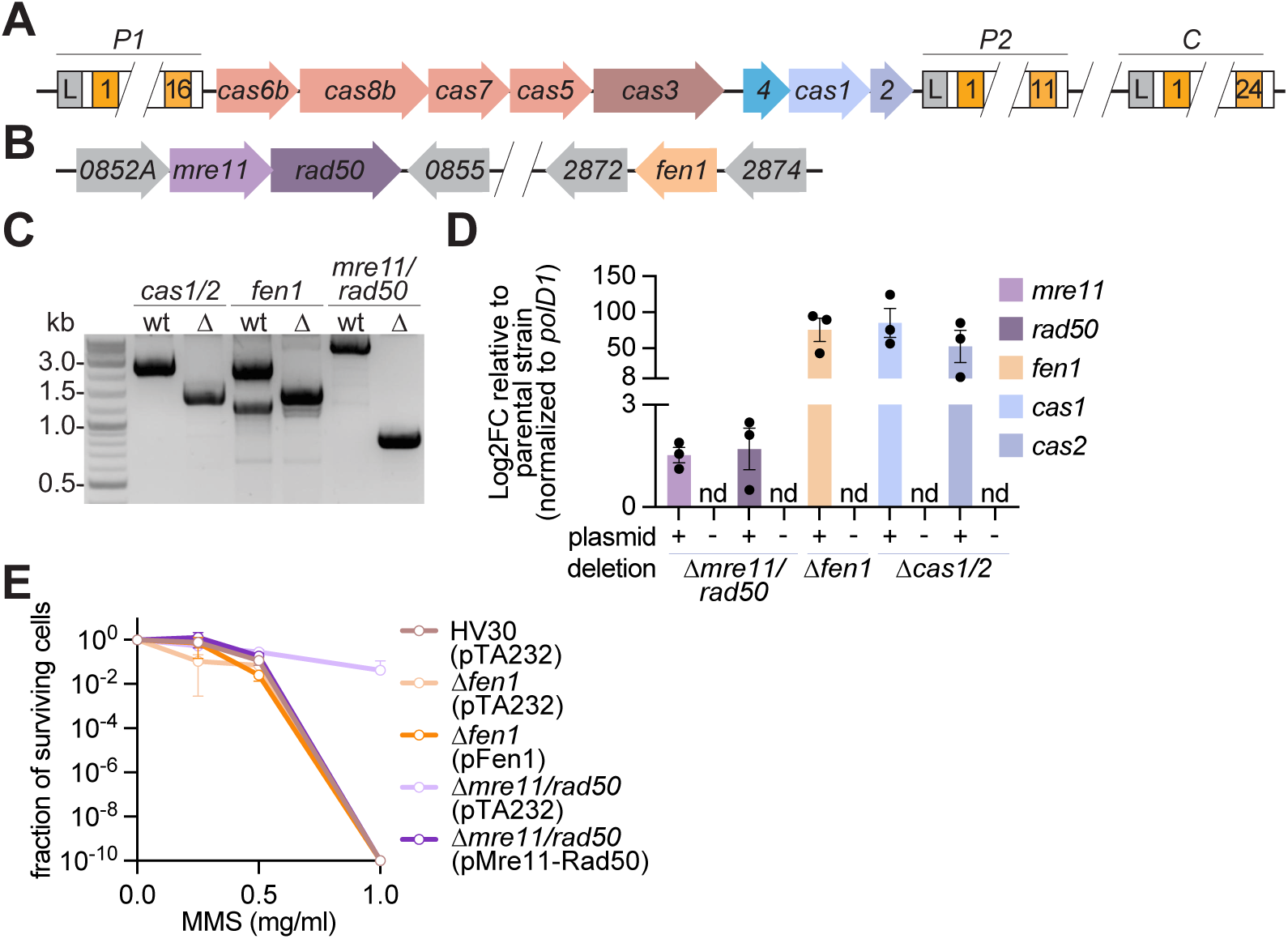
Generation of *mre11/rad50* and *fen1* deletion mutants in *H. volcanii*. **(A)** Schematic representation of the *H. volcanii* type I-B CRISPR-Cas system, found on the *H. volcanii* megaplasmid pHV4 (204.9 – 218.6 kb), and the C array on the main chromosome (2.385 – 2.387 Mb). **(B)** Schematic representation of the genomic regions in which *mre11/rad50* are found (0.764 – 0.769 Mb) and in which *fen1* is found (2.709 – 2.712 Mb) on the *H. volcanii* main chromosome. **(C)** Gel electrophoresis of PCR products spanning the deleted region in each mutant strain and in HV30. Expected product sizes are: 2706 bp (HV30) and 1448 bp (*Δcas1/2*) across *cas1/2* region using AC1071/AC1074, 2481 bp (HV30) and 1500 bp (*Δfen1*) across *fen1* region using AC495/AC498, and 4847 bp (HV30) and 847 bp (*Δmre11/rad50*) across *mre11/rad50* region using AC413/AC415. **(D)** RT-qPCR was used to find the Log2FC in gene expression of *mre11*, *rad50*, *fen1*, *cas1*, and *cas2* relative to parental strain HV30 and normalized to *polD1*. “nd” indicates not detected. **E)** *Δmre11/rad50* (pTA232) and *Δfen1* (pTA232) and their respective complementation strains, and HV30 (pTA232) were treated with methyl methanesulphonate (MMS) and incubated at 43°C for 1 hour before serial dilution and plating on Hv-YPC.

The DNA repair complexes RecBCD and AddAB present in Gram negative and positive bacteria, respectively, enhance spacer acquisition in Type I, II, III, and V systems (7–9, 30). This is because Cas1-Cas2 preferentially integrate sequences derived from free DNA ends into the CRISPR array, such as the injected end of a viral genome, and those that are the result of double stranded breaks (DSB) produced at chromosomal termini, restriction enzyme cut sites and Cas9 cleavage sites (4, 7, 8, 31). Free DNA ends are recognized and processed by RecBCD and AddAB to initiate repair by homologous recombination (HR) (32, 33). It has been hypothesized that this processing generates additional free DNA ends from which Cas1-Cas2 can acquire new spacers (7, 8). Since RecBCD/AddAB DNA degradation stops at *chi* sites, short regions of DNA that are highly enriched in the genome to prevent excessive degradation of self-DNA during repair (34), the region between a free DNA end and a *chi* site becomes a hotspot of spacer acquisition (7–9, 31). *Chi* sites are present in all bacteria studied to date but their sequences are highly variable (34, 35). HR also has affects DNA targeting by CRISPR-Cas systems. One study found that in *E. coli* cells infected with phage lambda, repair of the DNA cleaved by both type I and II CRISPR-Cas systems is efficiently repaired by the RecBCD-dependent HR pathway of the host (36). This repair is achieved with high fidelity and prevents the accumulation of mutations at the target site and the generation of “escape” phages that can bypass crRNA recognition. On the other hand, the viral recombination system, lambda-red, was found to significantly increase the frequency of escaper phages, presumably through the introduction of target mutations during viral-mediated HR repair (36).

Archaea have evolved different DNA repair proteins from those present in bacteria (37, 38), with many of them more closely related to the those found in eukaryotes (39). *H. volcanii* has been found to use both HR and microhomology-mediated end joining (MMEJ) to repair DSBs (40, 41), but it does not contain direct homologs of RecBCD or AddAB. DNA end resection during MMEJ and HR is initiated by Mre11-Rad50, a DNA repair complex conserved in bacteria (SbcCD) and eukaryotes (Mre11-Rad50 with additional components), capable of DNA binding, unwinding, and resectioning (42–45). The complex is made of a Mre11 dimer with dsDNA exonuclease and ssDNA endonuclease activities and two Rad50 subunits with DNA-binding activity (40, 42, 45). Powered by ATP hydrolysis, the Mre11-Rad50 complex scans DNA, detects breaks, cleaves broken DNA ends and hairpins, and degrades the 5’ strand, leaving a short 3’ strand overhang during the early stages of HR repair. Therefore, although not conserved at the amino acid or structural levels, the Mre11-Rad50 complex performs a similar function as RecBCD and AddAB in bacteria (42, 46, 47). In contrast to bacteria, however, the Mre11-Rad50 complex is not affected by *chi* sites in *H. volcanii,* which have not been identified on archaeal genomes (35). In addition, while this complex is essential for some archaea (48), deletion of these genes is tolerated in *H. volcanii*, where the *mre11/rad50* double mutant has been reported to show higher resistance but slower recovery to DNA damage than wild-type (49). A second DNA repair protein that is highly conserved in archaea and eukaryotes is flap endonuclease 1 (Fen1). Fen1 is involved in trimming 5’ ends during Okazaki fragment maturation, as well as playing a role in base excision repair (BER) and MMEJ (50, 51). It is not part of the HR pathway and its 5’ endonuclease activity is different from that of Mre11-Rad50 and RecBCD/AddAB. In *H. volcanii*, deletion of *fen1* is viable but leads to an increase in sensitivity to DNA damage and oxidative stress (52).

While bacteria and archaea share the same CRISPR-Cas systems, they have different DNA repair systems. Given the importance of RecBCD and AddAB in CRISPR spacer acquisition in bacteria, we wondered whether Mre11-Rad50 and/or Fen1 play a similar role in archaea, more specifically in the Type I-B CRISPR-Cas immune response of *H. volcanii*. We generated deletion mutants to compare the acquisition of new spacers and found that deletion of *mre11rad50*, but not of *fen1*, reduces the levels of spacer acquisition. We also investigated the effect of these deletions on the targeting of *H. volcanii* pleomorphic virus 1 (HFPV-1), a nonlytic virus which causes chronic infection of host cells (53). It was determined that none of the *Haloferax* repair systems affect viral targeting. Our results describe how CRISPR systems have adapted to provide anti-viral defense to hosts from different domains of life.

## RESULTS

### Generation of *mre11/rad50* and *fen1* deletion mutants in *H. volcanii*

To investigate the importance of DNA repair systems for CRISPR spacer acquisition we looked for the integration of spacer sequences from the host genome in the absence of infection, also known as “self-acquisition”. This approach overcomes the difficulties posed by the extremely low occurrence of spacer acquisition, which takes place in only one cell per 10^5^-10^6^ infected cells (5), and therefore dramatically reduces the number of cells available for spacer analysis. Self-acquisition, however, induces CRISPR targeting of the host genome (“self-targeting”), a highly genotoxic event that would eliminate cells that acquire new spacers from the population. To overcome this, previous studies that successfully studied self-acquisition spacer patterns and rates used a host strain with mutations in Cas proteins that prevent DNA destruction (7, 9, 36). Therefore, to study self-acquisition we used *H. volcanii* strain HV30 (54), a derivative of the H119 strain (Δ*pyrE2* Δ*trpA* Δ*leuB*) (55), which lacks the *cas3* nuclease and the *cas6b* crRNA-processing endonuclease genes from the type I-B CRISPR-*cas* operon present on the pHV4 megaplasmid (Fig. 1A).

The *mre11*/*rad50* operon and the *fen1* gene are located in different loci of the *H. volcanii* main chromosome (Fig. 1B). We used an established gene knockout system (54) to eliminate both of these genetic units and confirmed the deletion using PCR (Fig. 1C). For complementation studies, we cloned *mre11*/*rad50* and *fen1* into the pTA232 shuttle vector, consisting of a pHV2 origin of replication and a *leuB* gene for selection in minimal media lacking leucine (55), generating pMre11-Rad50 and pFen1, respectively. Complementation was corroborated via RT-qPCR, to calculate the expression of *mre11*, *rad50* and *fen1* normalized to *polD1* and relative to wild-type levels. The results showed that neither gene was detected in the respective deletion strains, that *mre11/rad50* mRNA levels are restored to wild-type levels when pMre11-Rad50 is present, and that *fen1* is over-expressed by pFen1 when compared with wild-type (Fig. 1D). We then tested the Δ*mre11/rad50* and *Δfen1* mutants’ response to DNA damage by the alkylating agent methyl methanesulfonate (MMS). Previous studies have found that after UV-C treatment, Δ*mre11/rad50* and Δ*fen1* mutants display decreased and increased sensitivity to DNA damage, respectively (49, 52). We observed an increase in survival for the Δ*mre11/rad50* deletion in the HV30 genetic background, but the Δ*fen1* mutant did not differ from wild-type when using this mutagen (Fig. 1E). The phenotype of the Δ*mre11/rad50* deletion was reverted to wild-type MMS sensitivity by the pMre11-Rad50 complementation plasmid (Fig. 1E).

In addition to the deletion of DNA repair genes, we also generated a strain lacking *cas1*/*cas2* (Fig. 1B), encoding for the spacer integrase (21), which we used as a control for the absence of spacer acquisition. This strain was complemented with a pTA232 plasmid carrying the deleted genes, pCas1-2. RT-qPCR corroborated the absence of *cas1* and *cas2* expression in the mutant, as well as the over-expression of these genes in the strain harboring pCas1-2 (Fig. 1D).

### *mre11/rad50*, but not *fen1,* are required for efficient spacer acquisition in *H. volcanii*

Previous studies have shown that the CRISPR-C array (Fig. 1A) is not active in spacer acquisition (10). We therefore initially investigated the addition of new spacers to the CRISPR-P1 and CRISPR-P2 arrays (Fig. 1A) of the wild-type and mutant HV30 strains. We developed a two-step, PCR-based method which involves an initial amplification of a large region upstream of the leader sequence (56, 57) and the first spacer sequence of the CRISPR array to eliminate all but the first repeat of the locus. This PCR product is used as template for a second amplification with primers that anneal within the leader sequence and the first repeat of the array (Fig. 2A). Finally, next generation sequencing of this smaller PCR product is used to find the sequence and number of reads of the newly acquired spacers. For each spacer whose sequence matches the HV30 genome, its abundance is normalized to the number of reads per million of total reads in the sequencing experiment to calculate the RPM value. To obtain the pattern of spacer distribution across the genome, we segmented the HV30 DNA sequence into 10 kb bins and calculated the average RPM values of spacers contained within each bin (36). The data for all next generation sequencing experiments carried out in this work is provided in the Supplementary Data file.

**Figure 2.**
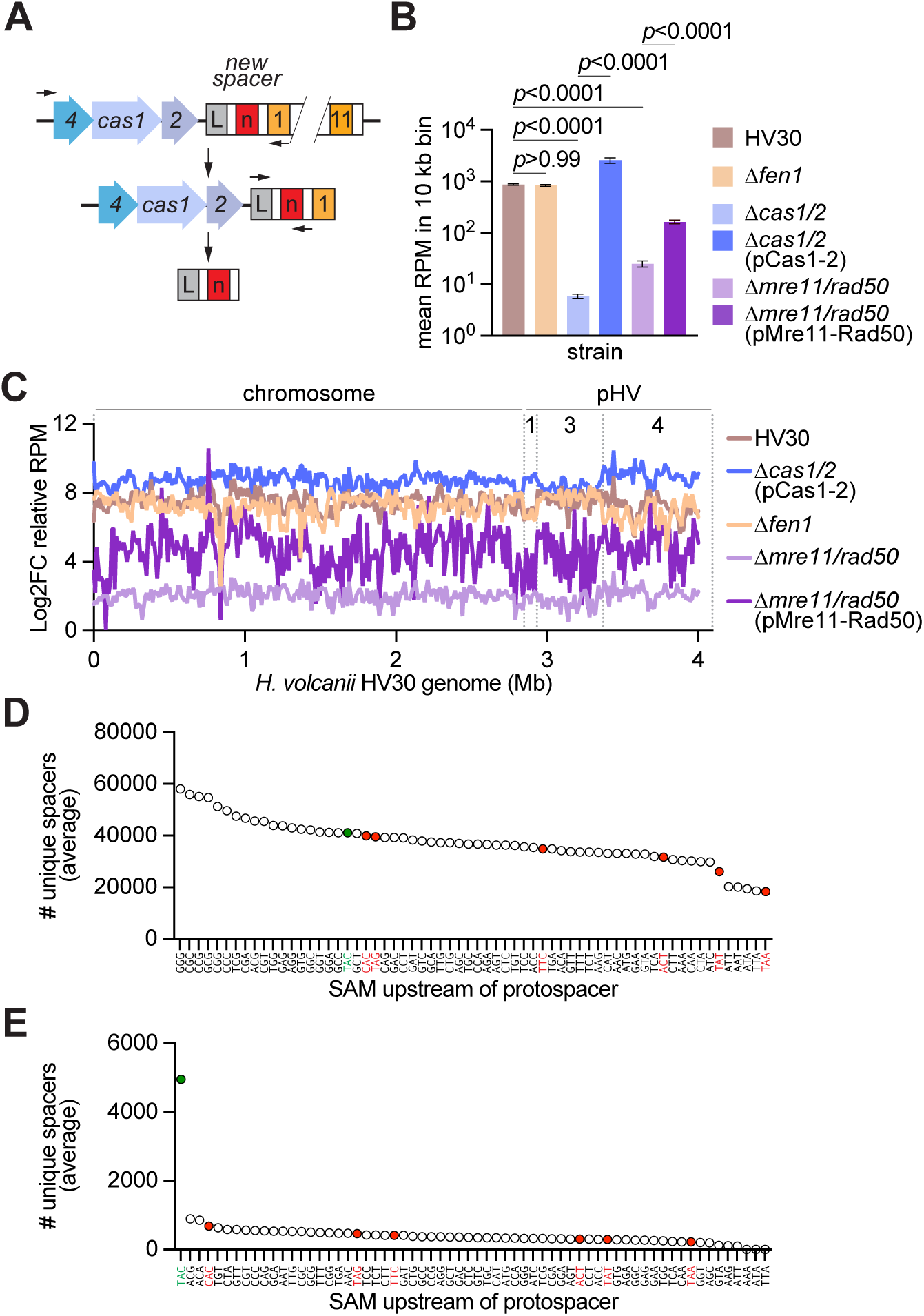
mre11/rad50, but not fen1, are required for efficient spacer acquisition in *H. volcanii.* **(A)** Two-step PCR design with primers (arrows) allowing for detection of rare acquisition events by the endogenous P2 array. **(B)** Average abundance (measured as reads per million of genome-matching reads (RPM) of spacers acquired into the P2 array per 10 kb bin across the *H. volcanii* genome. **(C)** Log2FC in acquired spacer abundance (n=3) (RPM) from *Δcas1/2* negative control, mapped to the archaeal genome (10 kb bins) (main chromosome, with megaplasmids pHV1, pHV3, and pHV4 denoted the dashed lines). **(D)** Average number of unique spacers with each SAM acquired into the P2 array in HV30, normalized for the frequency of the trinucleotide in the *H. volcanii* genome. **(E)** Average number of unique spacers with each SAM acquired into the P2 array in H119, normalized for the frequency of the trinucleotide in the *H. volcanii* genome. Green indicates the previously established primed self-acquisition SAM TAC, red indicates the previously established targeting PAMs.

We designed primers to adopt this strategy to evaluate self-acquisition of spacers into the CRISPR-P2 (Fig. 2A) and CRISPR-P1 arrays (Fig. S1A). We found that a much higher frequency of new spacers integrate into the P2 than into the P1 array, with the average RPM value for spacers acquired into the P2 array almost three orders of magnitude higher than the RPM value for P1 new spacers (Fig. S1B). The distribution of the P2 spacers across the HV30 genome was evenly increased, demonstrating that the new spacers did not originate from isolated hotspots of acquisition (Fig. S1C).

Therefore, we decided to evaluate the activity of this array in the Δ*mre11/rad50* and Δ*fen1* mutants, using the Δ*cas1/2* mutant as a control for the spacer acquisition noise. We found that the average RPM for the spacers acquired in the Δ*fen1* genetic background was similar to that of the HV30 parental strain (Fig. 2B). In contrast, the RPM values for spacers acquired in the Δ*mre11/rad50* mutant were significantly reduced to a similar extent as the values for the control strain that lacks the Cas1/2 spacer integrase (Fig. 2B). Restoration of Mre11-Rad50 expression from the complementing plasmid partially rescued spacer acquisition (Fig. 2B). Over-expression of Cas1/2, on the other hand, led to a marked increase in the abundance of new spacers (Fig. 2B), a result previously observed in bacteria, where increased production of the integrase led to markedly elevated rates of spacer acquisition in type I (57), II (5) and III (9) CRISPR-Cas systems. To visualize the spacer acquisition patterns of these strains we plotted the RPM values across the HV30 genome. We found an even distribution of values, without notable hot or cold spots, a result that demonstrates a general effect of these mutations on spacer acquisition (Fig. S1D). Nevertheless, peaks and valleys of small magnitude were observed in all patterns, including the negative control Δ*cas1/2* strain. Therefore, to eliminate false fluctuations we normalized the RPM values of each strain to the values of the control strain (Fig. 2C). The corrected patterns corroborated the previous results, showing that absence of Mre11 and Rad50, but not of Fen1, reduces spacer acquisition across the *Haloferax* genome.

### Type I-B CRISPR-Cas targeting affects the spacer acquisition motifs in *Haloferax*

Due to the key role of the PAM for targeting by CRISPR-Cas systems, several studies have explored whether all newly acquired spacer sequences have “spacer acquisition motifs” (SAMs) that match the PAM of the protospacer and therefore mediate efficient targeting (8, 28, 57). A previous study determined that the type I-B CRISPR-Cas system of *H. volcanii* can effectively target sequences downstream of TTC, ACT, TAA, TAT, TAG, or CAC motifs (58). In contrast to this PAM diversity, work that investigated primed self-acquisition revealed that the great majority of the new spacer sequences match a protospacer downstream of a SAM with the TAC sequence (10). The same study looked for SAMs during naïve self-adaptation but failed to find a consensus due to low number of sequences obtained. We therefore used our data to determine the 3-nucleotide upstream sequence of each new spacer and calculated the average frequency (among our three spacer acquisition replicas) for each of these SAMs, which was then corrected by the frequency of this trinucleotide in the HV30 genome. We found that in the absence of CRISPR targeting CGC, CCG, GCG and GCC motifs were somewhat preferred (Fig. 2D), even when we take into account the efficiency of acquisition by weighting in the number of reads of each spacer (Fig. S2A). Given that the previous study investigated primed adaptation using a self-targeting spacer, we wondered if our result would change when using the H119 strain which, as opposed to HV30, has functional *cas3* and *cas6b* genes and is capable of type I-B CRISPR targeting. We therefore collected next-generation sequencing data of spacer PCR products (Fig. 2A) from this strain and analyzed it to obtain the acquisition pattern. Compared to the non-targeting strain HV30, H119 self-acquisition displayed a marked reduction, of almost three orders of magnitude, in the integration of new spacers (Fig. S2B), and the distribution pattern was more fluctuating, most likely due to the low number of spacers that are averaged in each 10 kb bin (Fig. S2C). In most organisms, self-acquisition is reduced in the presence of an active CRISPR system due to the genotoxicity of the acquired self-targeting spacers (59–61). However, a previous study in *H. volcanii* showed that the introduction of a plasmid harboring a self-targeting spacer, which presumably would lead to the attack of the host chromosome and prevent the growth of transformants, was not impaired unless the *cas6b* gene was deleted (29). We wondered whether these previous results were particular to the spacer sequence selected for the study and therefore decided to introduce a different self-targeting spacer into the H119 strain. We constructed a version of the pTA232 plasmid (pSpcNT) harboring the constitutive promoter *p.fdx*, the leader sequence (57, 62) of the P1 CRISPR array, and two repeats flanking PaqCI restriction sites to clone new spacers generated by the annealing of complementary oligonucleotides with appropriate overhang sequences (Fig. S2D). To induce self-targeting we selected a protospacer sequence within the *HVO_2971* gene that is preceded by a TAA PAM used in previous studies (58) (Fig. S2E), and introduced the corresponding spacer sequence into pSpcNT, generating pSpc2971. While plating of the transformation reaction that introduced pSpcNT into strain H119 competent cells resulted in ∼100 colonies, no transformants were detected for pSpc2971 (Fig. S2F). Thus, our observations demonstrate that self-targeting in *Haloferax* can have genotoxic effects and therefore we interpret the marked reduction of acquired spacers in strain H119 compared to HV30 to be a consequence of auto-immunity in the presence of an active CRISPR system. We then analyzed the SAMs of the spacers acquired from strain H119 and found that TAC was markedly prevalent when spacers were considered irrespective of their number of reads (Fig. 2E), and TAA and TAC were the top two SAMs when reads for each sequence were weighted (Fig. S2F). Therefore, together with previous results (10), our data indicates that TAC is a preferred SAM of the type I-B CRISPR-Cas system of *H. volcanii*. In the absence of targeting (a condition generated in the laboratory), there is a greater diversity of SAM sequences.

### Genetic engineering of *Haloferax* virus HFPV-1

While self-acquisition provides a convenient method to compare spacer acquisition in different genetic backgrounds, CRISPR-Cas immunity has evolved to protect the host from viral and plasmid infections (2, 3). Therefore, we decided to investigate spacer acquisition of archaea infected with *Haloferax volcanii* pleomorphic virus 1 (HFPV-1), a chronic pleolipovirus that causes a small but noticeable growth inhibition of infected cells (53). To follow infection through the selection and visualization of hosts carrying this virus, we inserted the *pyrE2* and *gfp* genes into the intergenic region between *ORF9* and *ORF10*, each of them expressed under the constitutive ferredoxin promoter (*p.fdx*), generating HFPV-1* (Fig. 3A). *pyrE2* encodes the orotate phosphoribosyltransferase enzyme (OPRTase), which is essential for synthesizing uracil and therefore is a selectable genetic marker for the growth of *Haloferax* in media without this nucleotide (55). In addition, it can act as a counter-selection marker when cells are exposed to 5-Fluoroorotic acid (5-FOA), since OPRTase processes 5-FOA into 5-fluorouridine monophosphate (5-FUMP), a highly toxic metabolite (55). *gfp* was codon-optimized for *H. volcanii* and encodes for the green fluorescence protein (GFP) (63) and can be used to track infected cells via microscopy.

**Figure 3.**
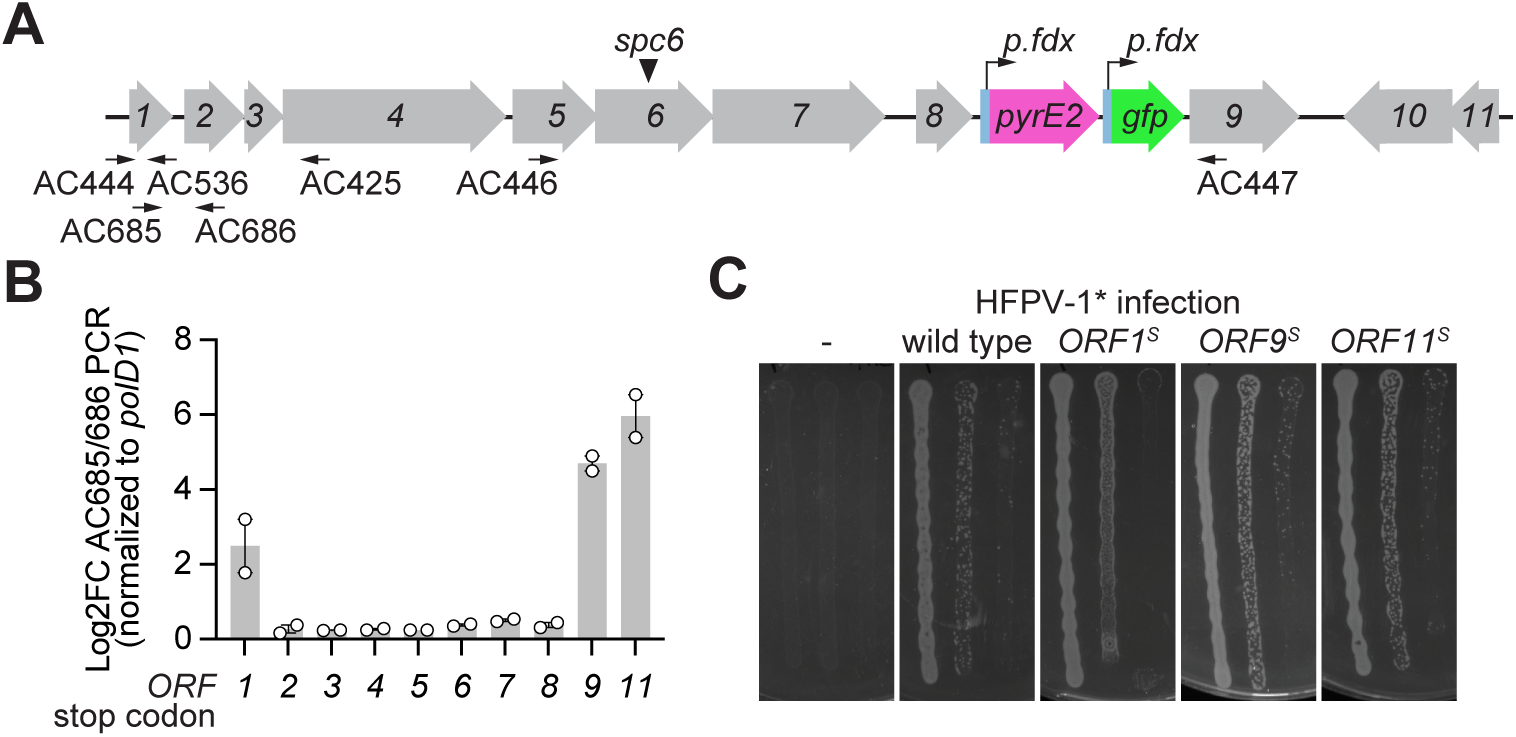
Genetic engineering of *Haloferax* virus (HFPV-1). **(A)** Schematic representation showing HFPV-1 genome (linearized) with *pyrE2* (pink) and *gfp* (green) engineered insertions, each under constitutive *p.fdx* promoters. **(B)** qPCR was used to measure the levels of viral DNA (using primers AC685/AC686) relative to the levels of genomic *polD1* DNA in the supernatants of H119 cultures infected with each of the HFPV-1 stop codons mutants. **(C)** Infectivity assay showing infectious viabilities of stop codon mutants ORF1^s^, ORF9^s^, and ORF11^s^ which showed elevated viral DNA in supernatants, with wild type HFPV-1* and uninfected controls.

We generated PCR products containing the inserted constructs as well as viral sequences, linked them using isothermal DNA assembly (55) to generate the engineered HFPV-1* genome, and introduced the virus into the parental strain H119 (Δ*pyrE2* Δ*trpA* Δ*leuB*) (55). Transformants were selected on plates containing minimal media lacking uracil. After confirming the sequence of the assembled mutant viruses, transformants were grown in liquid media and supernatants were collected to test for the presence of infectious particles by mixing with a fresh culture of H119 cells for 72 hours. Cells were plated and examined for the presence of HFPV-1* via PCR, which demonstrated that they contained the virus (Figs. 3A and S3A) and therefore were infected. Infections were also carried out using supernatants containing wild-type HFPV-1 and cells plated after 72 hours of growth in minimal media lacking uracil, where only cells incubated with HFPV-1*, but not HFPV-1, were able to form colonies (Fig. S3B). Inspection of cultures using fluorescence microscopy revealed green cells only after infection with HFPV-1* (Fig. S3C). Together, these results demonstrate that the engineered virus can infect and propagate in *H. volcanii*.

The success of our method to manipulate HFPV-1 using isothermal DNA assembly of PCR products prompted us to generate mutant viruses harboring stop codons in each of the 11 annotated viral ORFs, to investigate their importance in HFPV-1 replication and propagation. With the exception of the mutant lacking ORF10 expression, which encodes a putative replication protein (53), we obtained transformants that contained viral DNA, determined by PCR, for the mutants with disruptions in all the other ORFs (Fig. S3D). To test whether these mutants are capable of forming viral particles, we performed qPCR to quantify viral DNA in the supernatant of infected cultures 78 hours post infection. We found that mutations that prevent expression of ORF1, ORF9 and ORF11 were able to release particles containing viral DNA (Fig. 3B). Finally, to determine whether the particles are infectious we measured their ability to propagate into uninfected hosts. We mixed at a 1:1 ratio uninfected H119 cells carrying pTA232, a plasmid expressing *leuB* for selection in minimal media lacking leucine (55), with cells infected with each of the three ORF mutants as well as wild type HFPV-1* virus. Cells were collected after incubation on a membrane filter for 4 days and plated on minimal media lacking both leucine and uracil to enumerate colonies formed by newly infected hosts. We found that disruption of ORF1, ORF9 and ORF11 expression did not affect HFPV-1* propagation (Fig. 3C). These results describe a method to genetically engineer HFPV-1 and to generate a selectable version of this virus, which we employed to investigate its propagation as well as its interplay with the type I CRISPR-Cas system of *Haloferax volcanii*.

### Spacer acquisition during HFPV-1* infection

To measure spacer acquisition, we infected the parental HV30 strain as well as the Δ*mre11/rad50*, Δ*fen1* and Δ*cas1/2* mutants with HFPV-1*. We then used PCR amplification of the P2 CRISPR array and next generation sequencing to measure spacer acquisition. We also used uninfected cells as a control, which are only capable of self-acquisition given the non-targeting genetic background of the HV30 strain (Δ*cas3* Δ*cas6b*). When we compared the overall acquisition of this uninfected control to the values obtained for the parental HV30 strain carrying HFPV-1*, we observed a striking reduction in the detection of new spacers, ∼20 fold less in the infected strain (Fig. 4A). Moreover, the majority of the new spacers detected in the infected strains matched to the host genome; i.e., they were the result of self-acquisition (Fig. S3). Although reduced in their quantity, the effect of the deletions of DNA repair genes was similar to those observed in uninfected strains, with a reduction in acquisition for the Δ*mre11/rad50* and Δ*cas1/2* mutants, but not for the Δ*fen1* strain (Fig. 4A).

**Figure 4.**
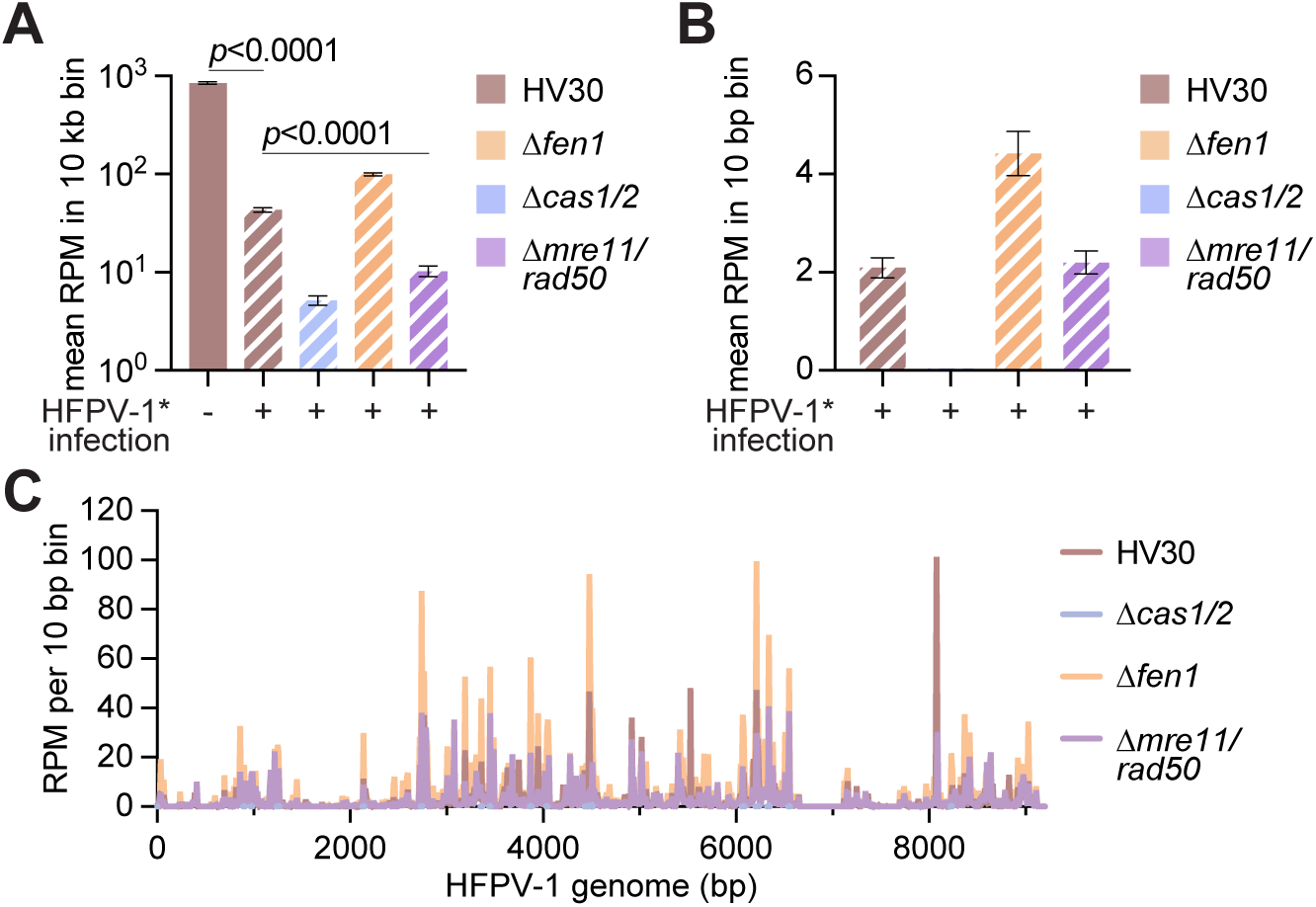
Spacer acquisition during HFPV-1* infection. **(A**) Average abundance (measured as reads per million of reads matching to genome or virus (RPM)) of spacers acquired into the P2 array per 10 kb bin across the *H. volcanii* genome. **(B)** Average abundance (RPM) of spacers acquired into the P2 array per 10 bp bin across the HFPV-1* genome. **(C)** Average abundance (RPM) of spacers acquired into the P2 array by the *H. volcanii* type I-B acquisition machinery, mapped to the HFPV-1* genome (10 bp bins).

For the small fraction of viral spacers that we were able to detect, however, we found that the Δ*cas1/2*, but not the Δ*mre11/rad50* deletion, reduced acquisition, and an increase in the number of spacers detected in the Δ*fen1* strain (Fig. 4B). The pattern of spacer acquisition (obtained by using 10 bp bins across the HFPV-1* virus) for the strains defective in DNA repair was similar to that of the parental HV30 strain (Fig. 4C). These results, however, are difficult to interpret given the very low numbers of spacer acquisition detected during infection. This decrease, on the other hand, is in line with previous work that reported the down-regulation of *cas* gene transcription in *Haloferax* cells infected with HFPV-1 (64), since high expression of the Cas1/2 integrase and Cas4 are required to obtain detectable levels of spacer acquisition in this organism (29). Overexpression of Cas1/2 is also required to increase the spacer acquisition rates in bacteria harboring type I (7), II (5) and III (9) CRISPR-Cas systems.

### *mre11/rad50* and *fen1* are not required for HFPV-1* type I-B CRISPR-Cas targeting

We also decided to test the effect of *mre11/rad50* and *fen1* deletions on targeting of HFPV-1*, since previous work showed that both bacterial and viral homologous-directed DNA repair systems affect the efficiency of type II-A targeting of lambda phage in *E. coli* (36). To do this, we generated the Δ*mre11/rad50* and Δ*fen1* deletions in the H119 strain (Fig. 5A), which has an intact type I-B CRISPR-Cas system and therefore is capable of targeting the virus. We selected a protospacer sequence within the *ORF6* gene (Fig. 3A) that is preceded by a TAT PAM used in previous studies(58), and introduced the corresponding spacer sequence into pSpcNT, generating pSpc6 (Fig. S2D). Both plasmids were transformed into the parental H119 strain and the Δ*mre11/rad50* and Δ*fen1* derivatives carrying HFPV-1*. Transformants were selected for growth in the absence of leucine (*leuB*, plasmid selection) and uracil (*pyrE2*, virus selection) and enumerated to calculate the transformation efficiency as colony forming units per µg of plasmid DNA. Compared with pSpcNT, the transformation efficiency of pSpc6 into H119 was three orders of magnitude lower (Fig. 5B), a result that demonstrates that the type I-B CRISPR-Cas system of *H. volcanii* can efficiently prevent the maintenance of the HFPV-1 virus. The deletions of the DNA repair genes, however, did not alter this difference. In addition, we incubated H119/pSpcNT and H119/pSpc6 archaea with cells harboring HFPV-1* (the same experiment described in Fig. 3C) to measure the ability of the type I-B CRISPR-Cas system to prevent viral infection. Cells were selected after this incubation for the presence of both *pyrE2* and *leuB* markers, i.e., plasmid-containing cells that were successfully infected with HFPV-1*. The number of colonies was normalized to the colonies obtained in plates that selected for the presence of plasmid only to calculate the efficiency of infection. Non-targeting conditions showed that approximately 1/1000 cells were infected by this method, however the presence of pSpc6 reduced viral transfer by 3-4 orders of magnitude (Fig. 5C). These results were unchanged in the mutant strains (Fig. 5C) and therefore we conclude that *mre11/rad50* and *fen1* are dispensable for CRISPR targeting.

**Figure 5.**
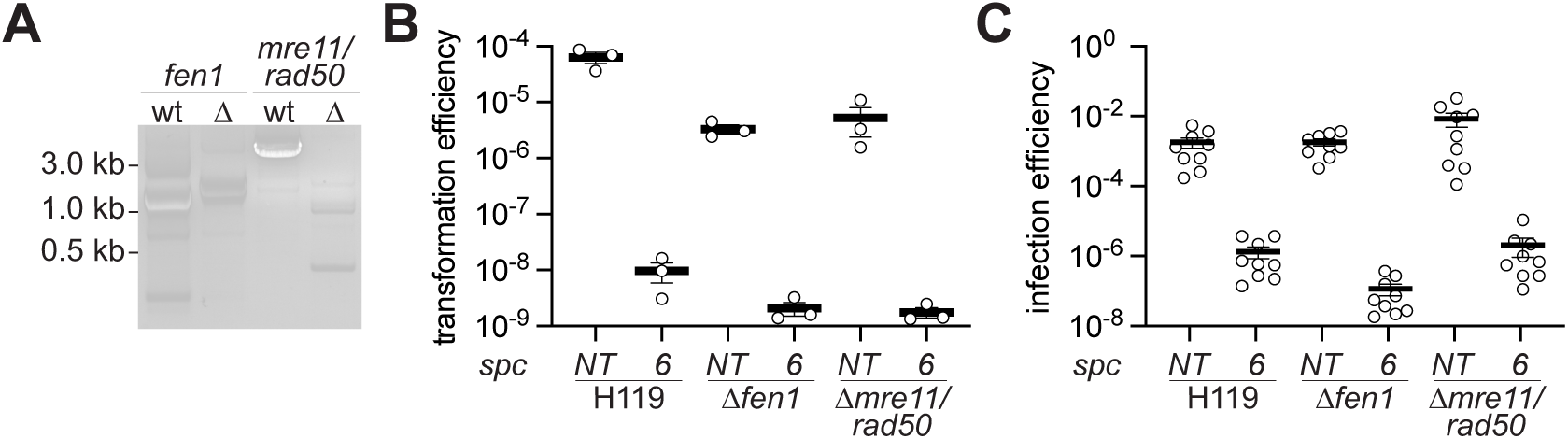
*mre11/rad50* and *fen1* are not required for HFPV-1* type I-B CRISPR-Cas targeting. **(A)** Gel electrophoresis of PCR products spanning the deleted region in each mutant strain and in H119. Expected product sizes are: 2481 bp (H119) and 1500 bp (*Δfen1*) across *fen1* region using AC495/AC498, and 4847 bp (H119) and 847 bp (*Δmre11/rad50*) across *mre11/rad50* region using AC413/AC415. **(B)** Transformation efficiencies of pSpcNT and pSpc6 into *H. volcanii* H119 strains infected with HFPV-1*. **(C)** Infection efficiencies using of HFPV-1* when infecting H119 strains containing pSpcNT or pSpc6.

## DISCUSSION

Here we studied the role of the DNA repair machinery in spacer acquisition and targeting during the Type I-B CRISPR-Cas response of *H. volcanii*. Our study showed that the Mre11-Rad50 complex is required, but not absolutely essential, for efficient spacer acquisition. In bacteria, the nucleases AddAB or RecBCD have a similar effect on spacer acquisition in type II-A (31, 36) and type I-E (7), and it is believed that the DNA degradation mediated by these complexes generates free DNA ends from which the Cas1-Cas2 integration complex can acquire new spacer sequences. It is not clear whether archaeal CRISPR-Cas systems also use free DNA ends as the source of new spacers. In bacteria, as a consequence of this mechanism, self-acquisition results in a high fraction of the new spacers being derived from a hotspot on the chromosomal terminus (7–9), where double-strand DNA breaks (DSBs) accumulate as a result of stalled replication forks at the termination (*ter*) sites (65). In this and a previous study in *H. volcanii* (10) however, such hotspots of spacer acquisition were not detected. This most likely is not due to the inability of Cas1/2 to use free DNA ends as substrates in archaea, but rather to differences in the mechanisms of DNA replication, which in archaea involves several origins of replication and lacks dedicated termination sites (66). The same previous study that looked for primed self-acquisition in the presence of chromosomal targeting found that at low levels of genotoxicity most spacers are derived from transposase genes, highly transcribed rRNA genes (both of which can display a high frequency of DSBs (9, 67, 68)), and the target site itself (also cleaved by the type I-B CRISPR-Cas system). Equivalent results were obtained in bacteria (8, 9). We did not detect particular hotspots, a result that may highlight differences between primed self-acquisition and the naïve self-acquisition experiments we performed. In addition, the previous study in *H. volcanii* showed that the over-expression of type IV restriction (*mrr*) or homing (*HEN*) endonucleases, also expected to generated numerous DSBs, did not increase the spacer acquisition frequency (10), a result that contrasts observations in bacteria where expression of the BglII restriction enzyme and of the I-Sce I endonuclease generated hotspots of spacer acquisition at the target sites (8, 9, 31). Our findings demonstrate that the Mre11-Rad50 complex, a functional homolog of RecBCD or AddAB with 5’-3’ DNA end resectioning activity, enhances spacer acquisition by the Type I-B system in *H. volcanii*, and therefore provide additional evidence that free DNA ends are important for the generation of new spacers in archaea. Last, we found that deletion of *fen1* did not affect spacer acquisition. Given that Fen1 is a flap endonuclease that acts on displaced RNA/DNA regions that accumulate during lagging strand replication (51), but does not process DSBs, this result also is in accordance with the role of free DNA ends as substrates for spacer acquisition.

Our extensive next-generation sequencing data allowed us to explore the presence of motifs flanking the protospacer sequence of the acquired spacers, known as the SAM. We found that in strain HV30, which lacks *cas6b* and *cas3*, CGC, CCG, GCG and GCC motifs were preferred. In contrast, self-targeting spacers acquired in strain H119, capable of DNA targeting, resulted in selection of the TAC motif. The same preference was observed in a study that looked at spacer acquisition in the presence of a self-targeting spacer (10). These previous experiments were performed in a *cas6b* deletion strain and therefore used an engineered system to transcribe a processed crRNA that attacked the host chromosome. Therefore, as opposed to our data that was obtained in the presence of a fully functional CRISPR system that can use new spacers to target the H119 genome, the new spacers acquired in the absence of the Cas6b endoribonuclease cannot produce mature, targeting crRNAs. In spite of this difference, both studies yielded the same SAM, which diverged from the SAM obtained in the HV30 strain in the absence of targeting. We believe that one possible factor that influences these results is the absence of Cas3 expression in HV30, which is present in the cells used in experiments that produced spacers with targets preceded by CGC, CCG, GCG and GCC SAMs. A role for Cas3 in the selection of sequences to be integrated as new spacers into the CRISPR array of the type I-E system of *E. coli* has been proposed after the finding that it associates with Cas1 in the adaptation complex (69). In addition, in type I-F systems Cas3 is a default component of the complex because it is naturally fused to Cas2 (70, 71). How Cas3 influences SAM sequence selection is not clear. Finally, from previously reported PAMs that license efficient targeting by the type I-B Cascade-Cas3 complex (TTC, ACT, TAA, TAT, TAG, or CAC (58)), only TAA was found to be a very strong SAM in our studies, an observation that reveals coordination between the immunization and targeting stages of the *H. volcanii* type I-B CRISPR response to ensure that a great majority of the acquired spacers are efficient at targeting.

In addition to investigating self-acquired spacer acquisition, we also studied spacer acquisition during infection with HFPV-1. Interestingly, the presence of the virus dramatically reduced spacer acquisition by a factor of 20. Therefore, we were not able to conclusively determine whether any of the DNA repair genes we deleted are important for CRISPR adaptation against this virus. We believe that this is a consequence of the previously reported downregulation of the type I-B *cas* genes, including *cas8*, *cas7*, and *cas1*, during chronic infection by this virus (64). The reduction of Cas1 expression will result in low concentrations of the Cas1/2 integrase, which has been determined to be detrimental for the detection of quantifiable levels of new spacers not only in *H. volcanii* (72), but also in the type I-E and type II-A CRISPR loci of *E. coli* and *S. pyogenes*, respectively (5, 57). We also tested the requirement of HFPV-1 genes for its maintenance and propagation and found that *ORF1*, *ORF9* and *ORF11* are dispensable for both. It is interesting to speculate that the non-essential role of one of these genes is to interfere with *cas* expression and/or limit spacer acquisition (73). Using AlphaFold3 (74) and Foldseek (75) we identified structural homologs of the ORF1 and ORF9, but not ORF11, proteins. Interestingly, ORF1 and ORF9 are putative transcriptional regulators with homology to the winged helix-like DNA-binding domains (76) of the MarR (77) and ArsR (78–80) families, respectively. HFPV-1 inhibition of the CRISPR-Cas response is specific to the immunization stage, since targeting was highly efficient and reduced the number of cells harboring the virus by four orders of magnitude. We believe that this could be explained by previous results obtained for the *E. coli* type I-E CRISPR-Cas system (as well as other) that showed that, as opposed to spacer acquisition that requires plasmid-borne over-expression of Cas1/2, targeting is less affected by the levels of the crRNA-guided Cas effector complexes. Therefore, targeting of, but not immunization with, HFPV-1 in *H. volcanii* may tolerate the inhibition on *cas* expression caused by the virus. Recently, it was reported that infection with another *Haloferax* virus (the lemon-shaped virus 48N; LSV-48N) reduces the *cas2*, *cas4*, *cas5* and *cas7* transcripts of the host (strain 48N) (81) and therefore it is possible that inhibition of CRISPR spacer acquisition via downregulation of *cas* genes is a common anti-defense strategy of haloarchaeal viruses. Alternatively, HFPV-1 may encode an inhibitor that specifically affects spacer acquisition, as it is the case for the truncated forms of the AcrIIA6 anti-CRISPR produced by *Streptococcus thermophilus* phage 123, which prevents spacer acquisition but not interference by the type II-A CRISPR-Cas systems of this organism (82). Future work will explore these possibilities as well as further investigate how the molecular differences between bacteria and archaea impact the function of CRISPR-Cas systems.

## MATERIALS AND METHODS

### Strains and growth conditions

Cultivation of *H. volcanii* strains was carried out in complete media (Hv-YPC), casamino acids media (Hv-Ca), or minimal media (Hv-Min)(83, 84). Liquid cultures were grown shaking (180 RPM) at 43°C and plates were grown at 45°C. Deletion strains in *H. volcanii* were constructed using the gene knockout system described previously(83). *E. coli* strains Turbo (NEB, Cat #C2984) and *dam-/dcm-* (NEB, Cat #2925) were grown in LB medium at 37°C, liquid cultures shaking (220 RPM), and supplemented with carbenicillin or ampicillin at 100 μg/mL as needed for plasmid maintenance. All strains, plasmids, and viruses used in this study are listed in Supplementary table 1.

### Plasmid construction

The plasmids used in this study are listed in Supplementary table 1. The oligonucleotide sequences used in this study are listed in Supplementary table 2. The plasmid cloning strategies used are listed in Supplementary table 3. Plasmids were transformed into *H. volcanii* using standard PEG-mediated transformation of haloarchaea (84).

### RT-qPCR

*H. volcanii* cultures were grown from picked colonies into Hv-YPC to OD_600_ 1.0. Cell pellets were collected from 1.5 mL of these cultures by centrifugation at 6800 rpm for 5 min. RNA was harvested from these pellets using an Agilent Absolutely Total RNA Purification Kit (Agilent, Cat #400800) with additional treatment of the RNA products with a Turbo DNA-*free* Kit (ThermoFisher, Cat #AM1907). Reverse transcription was carried out using 100 ng of template RNA from each sample with SuperScript IV Reverse Transcriptase (ThermoFisher, Cat #18090010) according to the manufacturer’s protocol. 40 ng of cDNA from each sample was used for qPCR using PowerUp SYBR Green Master Mix (ThermoFisher, Cat #A25777). qPCR was carried out with the QuantStudio 3 Real-Time PCR System (Applied Biosystems) set to Standard run mode and Standard Curve experiment type. The standard run cycle was 2 min at 50°C, 2 min at 95°C, 40 cycles of 1 sec at 95°C and 30 sec at 60°C, then a melt curve of 1 sec at 95°C, 20 sec at 60°C and an incremental increase from 60°C to 95°C at a rate of 0.15°C per sec. qPCR primers can be found in Supplementary table 2. Amount of RNA in each sample was calculated using the ΔΔCt method and was normalized using Ct values of housekeeping host gene PolD1 (HVO_0003).

Quantification of HFPV-1* viral shedding was carried out by qPCR of viral DNA extracted from supernatant 78 hours after infection. DNA was extracted from supernatant using PEG6000 as previously described (53). 40 ng of viral DNA extracted from supernatant was used for qPCR as described above.

### DNA damage assays

*H. volcanii* cultures were grown from picked colonies into Hv-YPC to OD_600_ 1.0. Cultures were divided into 250 µL aliquots and an appropriate amount of methyl methanesulfonate was added. Cultures were returned to 43°C shaking at 220 RPM for 1 hour. After incubation cultures were serially diluted in 18% salt water (84) and plated on Hv-YPC. Plates were incubated at 45°C for 5-6 days, until colonies were countable.

### *H. volcanii* gDNA extraction

*H. volcanii* cultures were grown from picked colonies into 3 mL Hv-YPC overnight, then were back-diluted 1:100 into 20 mL Hv-YPC, then back-diluted 1:100 into 500 mL Hv-YPC and grown to OD ∼1.0. Cell pellets were collected by centrifugation at 6800 rpm for 5 mins. Pellets were processed for gDNA extraction right away or kept at -80°C until ready to be processed. Genomic DNA extraction was carried out using a phenol-chloroform extraction protocol modified from an existing protocol (84). To a fresh or freshly thawed pellet, we add 10 mL of lysis solution (100 mM EDTA pH 8.0, 0.2% SDS) and 200 µL Proteinase K (ThermoFisher, Cat #EO0492) and vortexed vigorously. Lysates were incubated at 56°C for 10 min, vortexing occasionally, then treated with 40 µL RNase A (Qiagen, Cat #19101) and incubated at room temperature for 10 min. An equal volume of phenol:choloform:isoamyl alcohol (Millipore Sigma, Cat #P2069) was added to the lysate and vortexed for 30 sec. After centrifugation at 12000 rpm for 10 min at room temperature, the top aqueous layer was transfered to a fresh 50 mL Falcon tube. To each aqueous solution, we add 80 µL GlycoBlue coprecipitant (ThermoFisher, Cat #AM9515), 2.5 mL NH_4_OAc (7.5 M), and 18.75 mL cold 100% ethanol, and vortex. To precipitate DNA, tubes were placed at -20°C overnight. After overnight precipitation, tubes were centrifuged at 12000 rpm for 30 mins at 4°C. We then removed supernatants without disturbing the gDNA pellet and washed the pellets with cold 70% ethanol for two additional spins. Then pellets were dried and resuspended in 750 µL EB buffer (Qiagen, Cat #19086).

### Spacer acquisition and two-step PCR amplification of newly acquired spacers for next-generation sequencing

Genomic DNA extracted as described above was used as a template for PCR using a forward primer upstream of the array and a reverse primer on the first encoded spacer in the array. This PCR product was then amplified with a second PCR using primers at the end of the leader and in the repeat. Primers used in each of these PCR steps for amplification of P1 and P2 arrays can be found in Supplementary table 2. Preparation of the first PCR and the second PCR master mix before addition of the first round PCR product was conducted in a PCR-free cabinet to avoid contamination. Both PCR steps used Phusion High-Fidelity DNA polymerase (ThermoFisher, Cat #F534S). Cycling was performed with the following conditions: 98°C for 30 sec, 35 cycles of [98°C for 10 sec, annealing (see temperatures below) for 30 sec, 72°C for 15-30 sec/kb (see extension times below)], 72°C for 5 min, hold at 8°C. The first PCR step was 50 µL total volume per sample and used 1 µL gDNA as template, with additions of 10 µL betaine (Millipore Sigma, Cat #B0300) and 1.75 µL DMSO to improve amplification of from GC-rich template. First round amplification for the P1 array was annealed at 69°C with an extension time of 2 min, while the P2 array was annealed at 59°C with an extension time of 2 min, except for *Δcas1/2* strains which were had an extension time of 1 min. The round one PCR products were analyzed on 1.8% agarose gels stained with ethidium bromide and were extracted using a QIAquick Gel Extraction Kit (Qiagen, Cat #28704). The second round PCRs were set up in triplicate for each sample, with 250 ng of starting template each, which were pooled during PCR purification to increase the final amount of DNA in the sample. Second round amplification for both P1 and P2 arrays were annealed at 60°C with an extension time of 15 sec. PCR products were purified using a MinElute PCR Purification Kit (Qiagen, Cat #28004). Purified second round PCR products were size selected using E-Gel SizeSelect II Agarose Gels 2% with SYBR Gold II (ThermoFisher, Cat #GG661012) on the Invitrogen E-Gel Power Snap Electrophoresis Device (ThermoFisher).

These size selected products were prepared for sequencing using the TruSeq Nano DNA Library Prep protocol (Illumina, Cat #20015964). Final library molarities were determined with the 4200 TapeStation System (Agilent) and the Invitrogen Qubit 4 Fluorometer (ThermoFisher). Libraries underwent high-throughput sequencing with the MiSeq i100 Series (Illumina) using 25M 300 cycle reagent kits (Illumina, Cat #20126568) and a PhiX (Illumina, Cat #FC-110-3001) spike-in of 20%.

### Next-generation sequencing data analysis

Newly acquired spacers were extracted from MiSeq i100 fastq.gzip files using a Python code that finds sequences flanked by two repeat sequences. The spacers sequences, locations, reads, and abundance of all spacers in this study can be found in the Supplementary data file. Before mapping the spacers to the genome, we filtered the output to remove spacers that match to existing spacers in the P1, P2, or C arrays, as they may be the result of spacer rearrangement or artifacts of PCR, and not true newly acquired spacers (85, 86). We also filtered out spacers smaller than 23 bp or larger than 53 bp, as there is variation in spacer size centering on 36 bp, but spacers larger or smaller can be integrated(19) (Figure S1D). Spacers were mapped to bins across the *H. volcanii* genome or HFPV-1 genome using a Python code that sorts spacer reads or RPM by location into 10kb or 10 bp bins. RPM values were calculated as genomic reads (RPM_gen_) or genomic and viral reads (RPM_spc_) per million aligned reads.

The same Python code extracts the 10 base pairs upstream and 10 base pairs downstream of each spacer. Analysis of PAM/SAM was performed using this data. All custom Python scripts used for data analysis as well as the Supplementary data file are deposited at https://github.com/Marraffini-Lab/Cassel_etal_2026.

### Construction of HFPV-1* and HFPV-1 stop codon mutants

Wild type HFPV-1, HFPV-1(*pyrE2*), and HFPV-1(*pyrE2, gfp*), here abbreviated HFPV-1*, were constructed by Gibson assembly of gene fragments (FragmentGENE, Azenta Life Sciences). The gene encoding GFP was codon optimized for expression in *H. volcanii*. Gibson assembly of viral gene fragments was performed using NEBuilder HiFi DNA Assembly Master Mix (NEB, Cat #E2621). Gibson products were dialyzed for 45 min on 0.025 µm membrane filters (Millipore Sigma, Cat #VSWP01300). The assembled viral genomes were then transformed in competent *H. volcanii* cells using the standard PEG-mediated transformation method used for plasmids, with two additional washes with regeneration solution (Hv-YPC, 15% sucrose) (84).From cultures grown from cells carrying the transformed viral DNA, HFPV-1 viral preparations were isolated, purified, and used to infect new *H. volcanii* cultures as described previously (53).

HFPV-1 stop codon mutants were constructed by Gibson assembly of PCR products amplifying HFPV-1* DNA with primers inserting a stop codon into each viral ORF (Table S2). These PCR products were then assembled and transformed as described above. The cloning strategies used to construct these viral mutants can be found in Supplementary table 3.

### HFPV-1* fluorescence microscopy

Turbid cultures of stationary phase *H. volcanii* cells (pink-red color, OD_600_ ∼1.2-2.0) either uninfected or infected with HFPV-1* were diluted 1:100 in Hv-YPC. We use microfluidics plates designed for bacterial use (Millipore Sigma, #Cat B04A-03) with the CellASIC ONIX2 system. Before loading cells into the microfluidics chambers, flow 18% salt water through all wells that *H. volcanii* cells will come in contact with to flush out any traces of PBS that may lyse cells. After loading, cells are trapped in the chamber and Hv-YPC media is flowed over them at a 5 µL/hour flow rate.

Phase contrast images were captured every 5 min for 36 hours while GFP signal was captured every 30 min at 5% power and 500 ms exposure time. Cells were maintained at 43°C using a Tokai HIT thermal box (Zeiss). Imaging was performed with a Nikon Eclipse Ti2e inverted microscope using a CFI60 Plan Apochromat Lambda Phase Contrast DM 100x Oil Immersion objective lens (Nikon) with a Zyla 4.2 sCMOS (Andor) camera (65 nm pixels) and an X-Cite Xylis LED Illuminator (Excelitas).

### HFPV-1* infectivity assays

We developed an assay for testing infectious viability of HFPV-1* modified from conjugation assays, based on the concept that there may be greater infectivity when cells are in direct contact. Uninfected cells (carrying a *leuB* marked plasmid) to infected cells (carrying *pyrE2* marker) were grown in Hv-YPC to late exponential/early stationary phase (OD_600_ ∼0.8-1.2). The uninfected and infected cells were normalized to the same OD, after which they were combined 1:1 (100 µL each) in 10 mL of Hv-YPC and concentrated onto a 0.45 µm membrane filter (Millipore Sigma, Cat #HAWP04700) using vacuum filtration. Filter discs were placed on Hv-YPC plates and incubated at 45°C for 4 days. The cells grown on the filter discs were then resuspended in 2 mL Hv-YPC by vortexing and serial dilutions of the resuspension were plated on i) Hv-YPC (all cells positive control), ii) Hv-Min with uracil and tryptophan (cells containing *leuB*), and iii) Hv-Min with tryptophan (cells containing *leuB* and *pyrE2*).

### CRISPR-Cas targeting assay

To test CRISPR-Cas targeting of the HFPV-1* virus, competent *H. volcanii* cells infected with HFPV-1* were transformed with pSpcNT (plasmid expressing a non-targeting spacer with PaqCI cut sites) or pSpc6 (plasmid expressing a 35 bp long spacer targeting in ORF6 of HFPV-1 with a TAT PAM). Serial dilutions of transformed cells were plated to determine transformation efficiencies therefore targeting efficiency by plating on Hv-Ca with tryptophan (cells that contain *pyrE2*) and on Hv-Min with tryptophan (cells that contain *leuB* and *pyrE2*).

To test CRISPR-Cas self-targeting of the host chromosome, competent H. volcanii cells were transformed with pSpcNT or pSpc2971 (plasmid expressing a 37 bp long spacer targeting in HVO_2971 with a TAA PAM). Serial dilutions of transformed cells were plated to determine transformation efficiencies and therefore targeting efficiency by plating on Hv-YPC (all cells) and on Hv-Min with uracil and tryptophan (cells containing leuB).

## Acknowledgements

We thank Anita Marchfelder at Ulm University for the *H. volcanii* strains H119 and HV30, and plasmid pTA232. We thank Heather Schiller and Mechthild Pohlschröder at the University of Pennsylvania for plasmid pTA131. We would like to thank past and present members of the Marraffini lab for their thoughtful discussions and advice. AKC was supported by the Philip Levine M.D. Graduate Fellowship at The Rockefeller University. LAM is an investigator of the Howard Hughes Medical Institute.

## Author contributions

AKC and LAM designed and conceived the study. AKC performed all experiments, with help from HC on HFPV-1 engineering and qPCR. AKC and LAM wrote the manuscript. All authors read and approved the manuscript.

## Competing interests

LAM is a cofounder and Scientific Advisory Board member of Intellia Therapeutics, a cofounder of Eligo Biosciences and a Scientific Advisory Board member of Ancilia Biosciences.

---

**Supplementary Figure S1.**
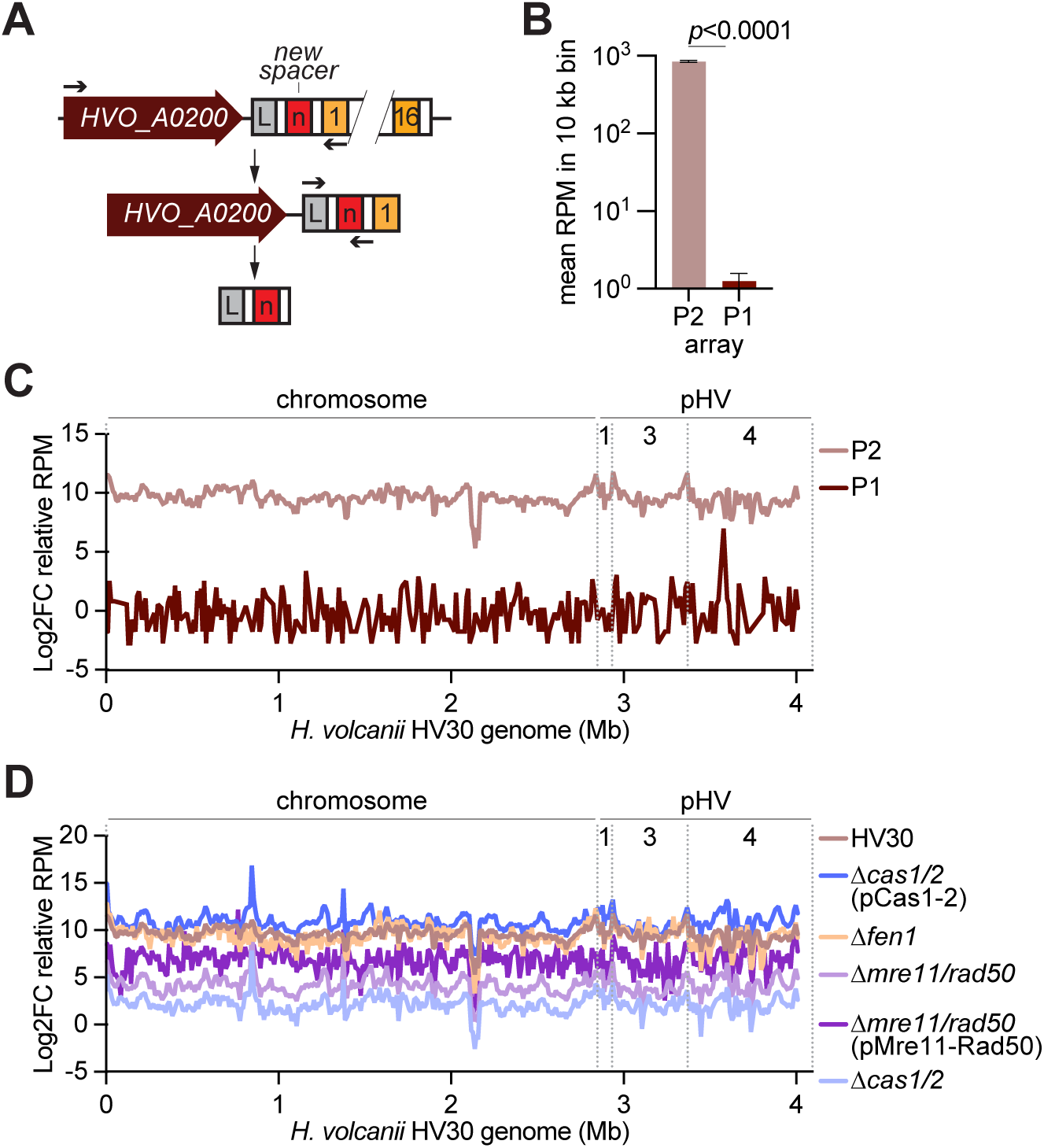
Spacer acquisition by the *H. volcanii* type I-B CRISPR-Cas system. **(A)** Two-step PCR design with primers (arrows) allowing for detection of rare acquisition events by the endogenous P1 array. **(B)** Average abundance (measured as reads per million of genome-matching reads (RPM) of spacers acquired into the P1 array per 10 kb bin across the *H. volcanii* genome. **(C)** Log2FC relative acquired spacer abundance (RPM) into P1 and P2 arrays, mapped to the archaeal genome (10 kb bins). **(C)** Log2FC relative acquired spacer abundance (RPM) of HV30, *Δcas1/2*, *Δfen1*, *Δmre11/rad50*, and their respective complementation strains mapped to the archaeal genome (10 kb bins) (main chromosome, with megaplasmids pHV1, pHV3, and pHV4 denoted the dashed lines).

**Supplementary Figure S2.**
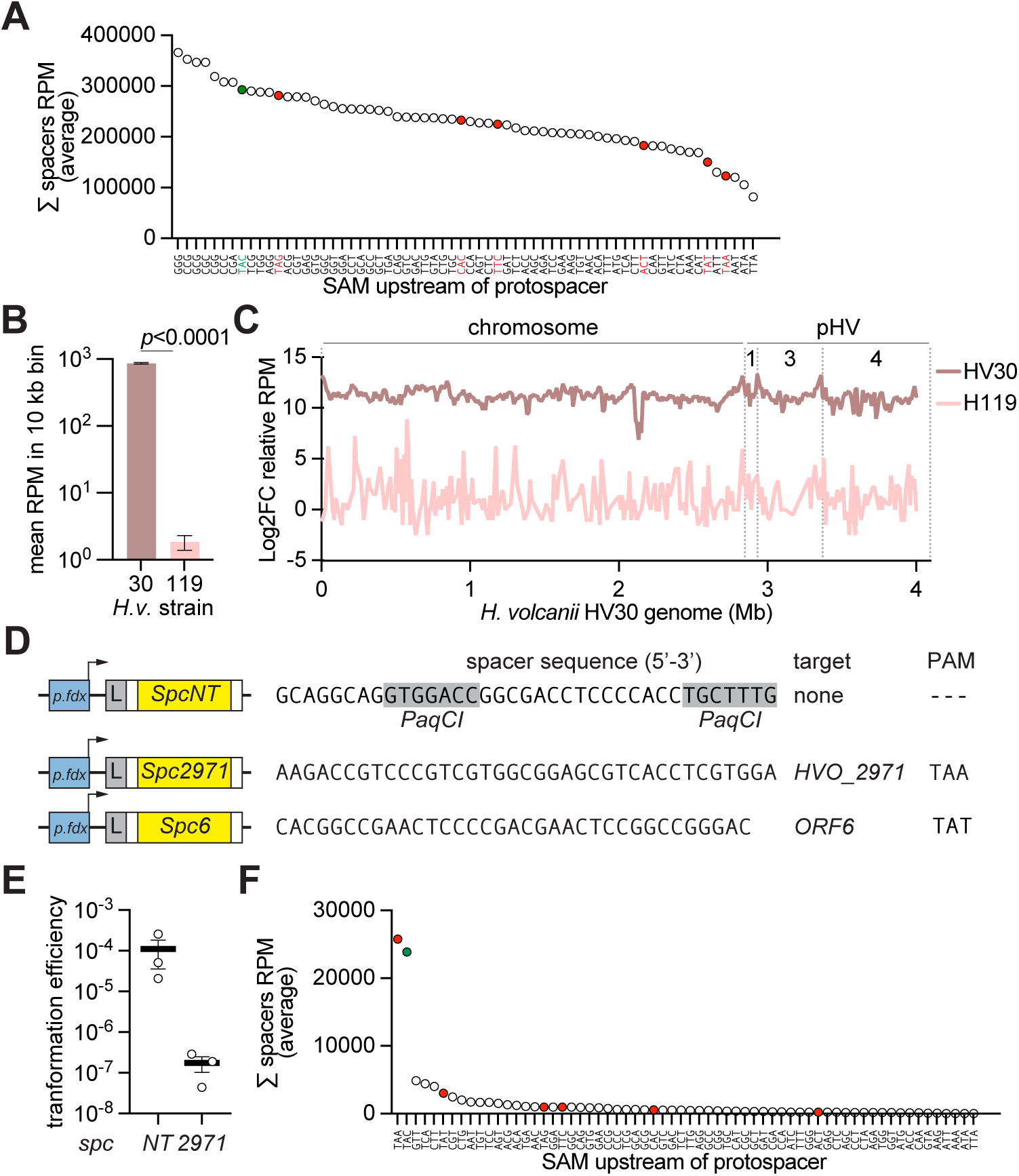
Self-targeting spacer acquisition by the *H. volcanii* type I-B CRISPR-Cas system. **(A)** Average sum of RPMs of unique spacers with each SAM acquired into the P2 array in HV30, normalized for the frequency of the trinucleotide in the *H. volcanii* genome. Green indicates the previously established primed self-acquisition SAM TAC, red indicates the previously established targeting PAMs. **(B)** Average abundance (measured as reads per million of genome-matching reads (RPM) of spacers acquired into the P2 array in HV30 and H119 per 10 kb bin across the *H. volcanii* genome. **(C)** Log2FC relative acquired spacer abundance (RPM) into the P2 array in HV30 and H119, mapped to the archaeal genome (10 kb bins) (main chromosome, with megaplasmids pHV1, pHV3, and pHV4 denoted the dashed lines). **(D)** Spacer sequences of each plasmid: pScpNT (non-targeting with PaqCI cut sites), pSpc2971 (self-targeting with TAA PAM), and pSpc6 (HFPV-1* targeting with TAT PAM). **(E)** Transformation efficiencies of pSpcNT and pSpc2971 into *H. volcanii* H119. **(F)** Average sum of RPMs of unique spacers with each SAM acquired into the P2 array in H119, normalized for the frequency of the trinucleotide in the *H. volcanii* genome. Green indicates the previously established primed self-acquisition SAM TAC, red indicates the previously established targeting PAMs.

**Supplementary Figure S3.**
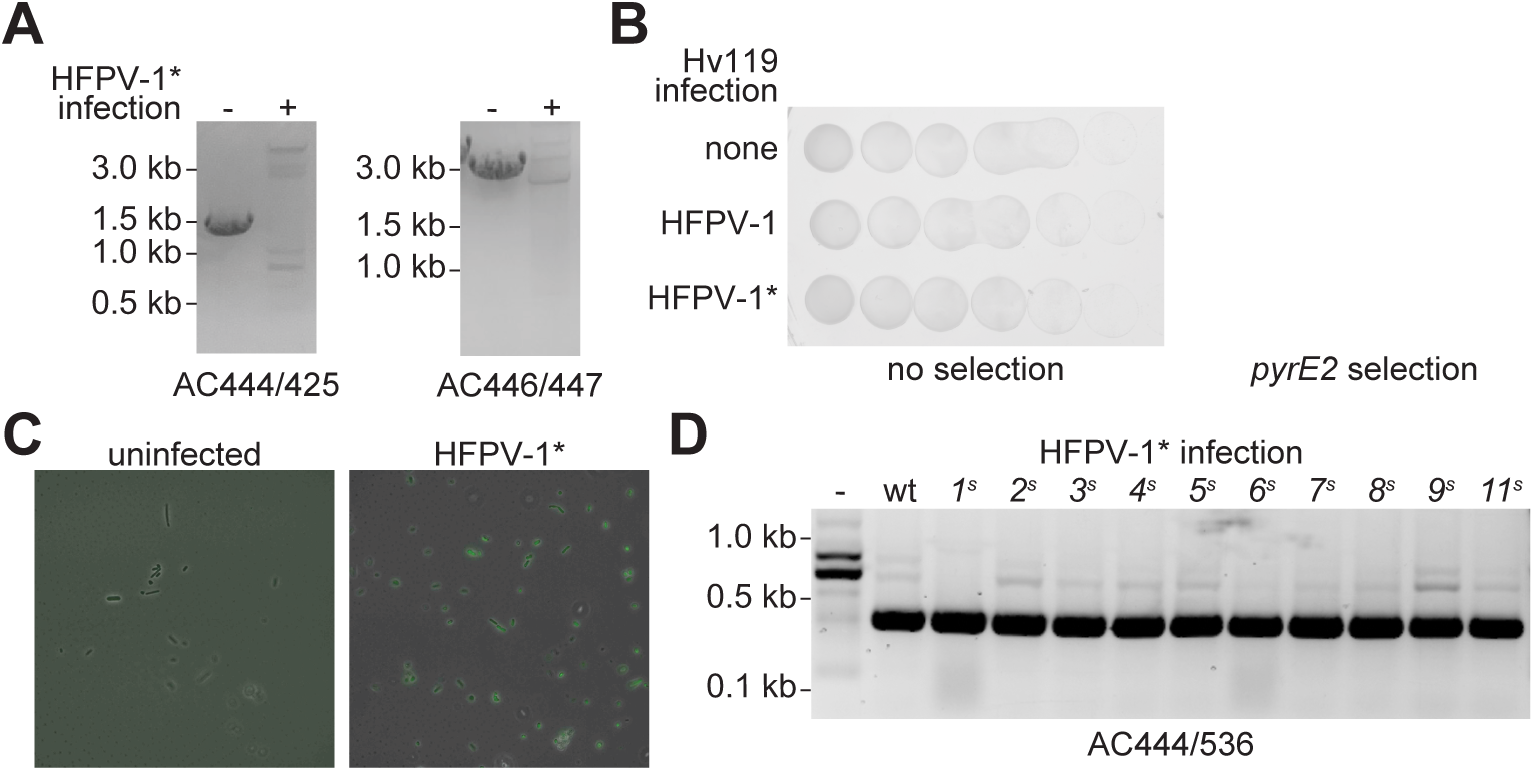
Genetic engineering of *Haloferax* virus (HFPV-1). **(A**) Gel electrophoresis of PCR products of gDNA extracted from cells infected with HFPV-1*. Expected product sizes are: 1207 bp using AC444/AC425 and 3000 bp using AC446/AC447. **(B)** Uninfected H119, H119 infected with WT HFPV-1, and H119 infected with HFPV-1* were plated on rich media with no selection and *pyrE2* selective media for cells carrying. **(C)** Microscopy showing GFP expression in cells infected with HFPV-1*. **(D)** Gel electrophoresis of PCR products of gDNA extracted from cells infected with HFPV-1* stop codon mutants. Expected product size is 406 bp using AC444/AC536.

**Supplementary Figure S4.**
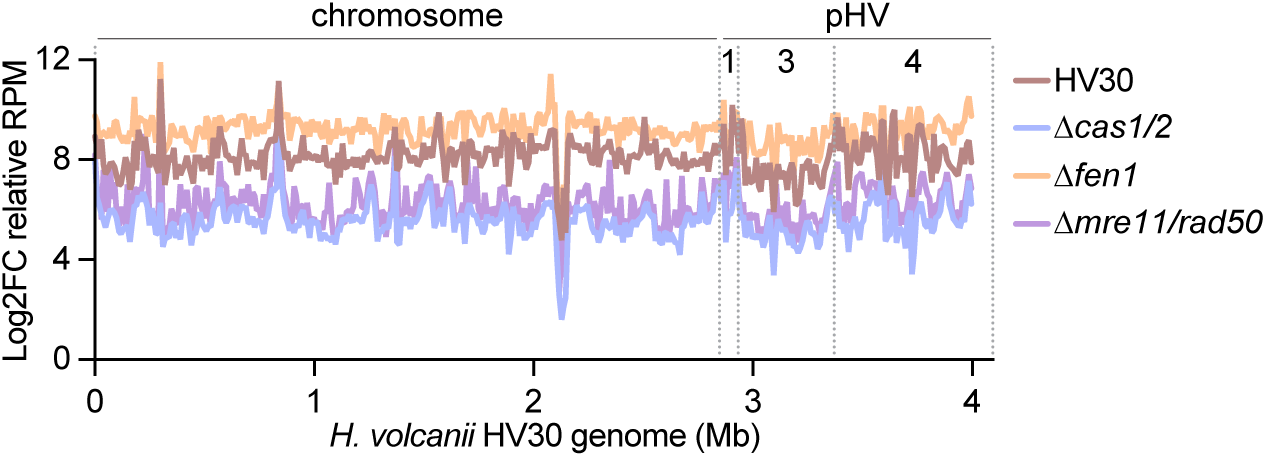
Spacer acquisition during HFPV-1* infection. **(A)** Log2FC relative acquired spacer abundance (RPM) into the P2 array of strains infected with HFPV-1* per 10 kb bin across the *H. volcanii* genome (main chromosome, with megaplasmids pHV1, pHV3, and pHV4 denoted the dashed lines).

## Notes

https://github.com/Marraffini-Lab/Cassel_etal_2026

