## Supplementary Tables for "Mre11-Rad50 enhances spacer acquisition in a haloarchaeal Type I-B CRISPR-Cas system"

**Supplementary table 1.** Strains, plasmids, and viruses used in this study.

| ***H. volcanii* strain** | **Description** | **Reference** |
| --- | --- | --- |
| H119 | DS70 *∆pyrE2 ∆leuB ∆trpA* | (1) |
| HV30 | DS70 *∆pyrE2 ∆leuB ∆trpA ∆bgaH ∆cas6 ∆cas3* | (2) |
| HVAC01 | HV30 *∆mre11/rad50* | This study |
| HVAC02 | HV30 *∆fen1* | This study |
| HVAC03 | HV30 *∆cas1/2* | This study |
| HVAC04 | H119 *∆mre11/rad50* | This study |
| HVAC05 | H119 *∆fen1* | This study |
| HVAC06 | H119 *∆cas1/2* | This study |
| ***E. coli* strain** | **Description** | **Reference** |
| Turbo | cloning and plasmid storage strain | New England Biolabs  Cat #C2984 |
| dam-/dcm- | methyltransferase deficient *E. coli* for passage of *H. volcanii* shuttle vectors | New England Biolabs  Cat #2925 |
| **Plasmid name** | **Description** | **Reference** |
| pTA131 | pBluescript II with *Bam*HI-*Xba*I fragment containing *pyrE2* under *p.fdx* promoter | (1) |
| pTA232 | pBluescript II with *leuB* under *p.fdx* promoter and *Nco*I-*Hind*III fragment containing pHV2 replication origin | (1) |
| pAC04 | pTA131-based integrative plasmid for with flanking regions for deletion of *mre11/rad50* | This study |
| pAC27 | pTA131-based integrative plasmid for with flanking regions for deletion of *fen1* | This study |
| pAC83 | pTA131-based integrative plasmid for with flanking regions for deletion of *cas1/2* | This study |
| pAC40 | pTA232-based shuttle vector with *p.fdx* promoter | This study |
| pMre11-Rad50 (pAC95) | pTA232-based shuttle vector expressing *mre11/rad50* under its natural promoter | This study |
| pFen1 (pAC42) | pTA232-based shuttle vector expressing *fen1* under *p.fdx* | This study |
| pCas1-2 (pAC86) | pTA232-based shuttle vector expressing *cas1/2* under *p.fdx* | This study |
| pAC23 | pTA232-based shuttle vector with P1 leader, repeat, original first genomic spacer, repeat | This study |
| pAC28 | pTA232-based shuttle vector with P1 leader, repeat, one PaqCI cut site in place of spacer, repeat | This study |
| pAC29 | pTA232-based shuttle vector with P1 leader, repeat, twp PaqCI cut sites in place of spacer, repeat | This study |
| pSpcNT (pAC53) | pTA232-based shuttle vector with *p.fdx* expressing: P1 leader, repeat, PaqCI cut sites in place of spacer, repeat | This study |
| pSpc6 (pAC70) | pTA232-based shuttle vector with *p.fdx* expressing: P1 leader, repeat, 35 bp spacer targeting HFPV-1 ORF6 with TAT PAM, repeat | This study |
| pSpc2971 (pAC59) | pTA232-based shuttle vector with *p.fdx* expressing: P1 leader, repeat, 37 bp spacer targeting HVO_2971, with TAA PAM, repeat | This study |
| **Virus** | **Description** | **Reference** |
| HFPV-1 | Haloferax pleomorphic virus 1 | (3) |
| HFPV-1 *(pyrE2)* | HFPV-1 engineered with *pyrE2* expressed under a *p.fdx* promoter inserted after *ORF9* | This study |
| HFPV-1* | HFPV-1 engineered with *pyrE2* and *gfp* each expressed under *p.fdx* promoters inserted after *ORF9* | This study |
| HFPV-1*-ORF1^s^ | HFPV-1*with stop codon in *ORF1* | This study |
| HFPV-1*-ORF2^s^ | HFPV-1* with stop codon in *ORF2* | This study |
| HFPV-1*-ORF3^s^ | HFPV-1* with stop codon in *ORF3* | This study |
| HFPV-1*-ORF4^s^ | HFPV-1* with stop codon in *ORF4* | This study |
| HFPV-1*-ORF5^s^ | HFPV-1* with stop codon in *ORF5* | This study |
| HFPV-1*-ORF6^s^ | HFPV-1* with stop codon in *ORF6* | This study |
| HFPV-1*-ORF7^s^ | HFPV-1* with stop codon in *ORF7* | This study |
| HFPV-1*-ORF8^s^ | HFPV-1* with stop codon in *ORF8* | This study |
| HFPV-1*-ORF9^s^ | HFPV-1* with stop codon in *ORF9* | This study |
| HFPV-1*-ORF11^s^ | HFPV-1* with stop codon in *ORF11* | This study |

**Supplementary table 2.** Oligonucleotides used in this study.

| **Name** | **Sequence (5’ to 3’)** | **Purpose** |
| --- | --- | --- |
| AC405 | AGCAATGCTATTAGATAATATTTGCTATCCGGATCCACTAGTTCTAGAGCG | pAC04 cloning |
| AC406 | GCATCGGCCCCACCGTGAGTCCCGCGAGGCCCCGGGCTGCAGGAATTC | pAC04 cloning |
| AC401 | GCCTCGCGGGACTCACGGTG | pAC04 cloning |
| AC407 | GGGCGGTCGGCTCCGCCG | pAC04 cloning |
| AC408 | AGTAGGCTATCGCGGCGGAGCCGACCGCCCCTATCACTCGGTTCGCCAC | pAC04 cloning |
| AC409 | GGATAGCAAATATTATCTAATAGCATTG | pAC04 cloning |
| AC493 | CGTTCGGTTCTTCGACCGCTTCGCACCCGTTGCAGCCCGGGGGATCCAC | pAC27 cloning |
| AC494 | TCCGTCGGCGGCGACCCCGACTCGGTAGACGGAATTCGATATCAAGCTTATCGATACCGT | pAC27 cloning |
| AC495 | GTCTACCGAGTCGGGGTCG | pAC27 cloning, *fen1* spanning on genome |
| AC496 | GGCGCGCTCGCATCGAGAC | pAC27 cloning |
| AC497 | GGCGGGCTGTTGTCTCGATGCGAGCGCGCCTTGCTCGGCTTTCGGGGG | pAC27 cloning |
| AC498 | ACGGGTGCGAAGCGGTCG | pAC27 cloning, *fen1* spanning on genome |
| AC1069 | GGAACAAGCAATAGAGGACTTTGAAGGATGGGATCCACTAGTTCTAGAGCG | pAC83 cloning |
| AC1070 | GCTTCGTGACTCGATGTGGTGAGGAATTCTCCCGGGCTGCAGGAATTC | pAC83 cloning |
| AC1071 | AGAATTCCTCACCACATCG | pAC83 cloning, *cas1/2* spanning on genome |
| AC1072 | GGGTCACATCCAACAGAG | pAC83 cloning |
| AC1073 | CTGTACCAAGACCTCTGTTGGATGTGACCCACTAAGGCACCACCTTGAC | pAC83 cloning |
| AC1074 | CATCCTTCAAAGTCCTCTATTG | pAC83 cloning, *cas1/2* spanning on genome |
| AC1135 | CGCCCGCAGTAATTCGACTCGTCGGGAAACCTGTCGTG | pAC95 cloning |
| AC1175 | CATCCGGCATAGGGTGCAG | pAC95 cloning |
| AC1176 | CCTGCACCCTATGCCGGATGTGGAAAGCGGGCAGTGAG | pAC95 cloning |
| AC1138 | GAGTCGAATTACTGCGGGC | pAC95 cloning |
| AC589 | GACCGCTGGACGTAGGGCCGCCACCGCGGTGGA | pAC42 cloning |
| AC590 | CACTAGTTCTAGAGCATGGGAAACGCAGACCTGC | pAC42 cloning |
| AC591 | ACCGCGGTGGCGGCCCTACGTCCAGCGGTCGAGG | pAC42 cloning |
| AC592 | GTCTGCGTTTCCCATGCTCTAGAACTAGTGGATCCCCCGGGCTG | pAC42 cloning |
| AC1083 | CGGGGAGTCGGTTTACCTGAGGCCGCCACCGCGGTGGA | pAC86 cloning |
| AC1084 | TGGTGATTTGCTTTTGTCATGCTCTAGAACTAGTGGATCCCCCGGGC | pAC86 cloning |
| AC1085 | ATGACAAAAGCAAATCACCATATC | pAC86 cloning |
| AC1086 | TCAGGTAAACCGACTCCC | pAC86 cloning |
| AC601 | GAATTCGATATCAAGTGCAGAGTTCGGCTTCCGAACGCATAGTAGTTGCTG | pAC40 cloning |
| AC602 | GACGGTATCGATAAGCAGCAACTACTATGCGTTCGGAAGCCGAACTCTGCA | pAC40 cloning |
| AC603 | GCATAGTAGTTGCTGCTTATCGATACCGTCGACCTC | pAC40 cloning |
| AC604 | AAGCCGAACTCTGCACTTGATATCGAATTCCTGCAGC | pAC40 cloning |
| AC452 | CGCCCGCCGGCTCCTACG | pAC23 cloning |
| AC486 | CGTGTAACCGTCTTCTTTGATGGACTTGTGCA | pAC23 cloning |
| AC482 | TTGTGGGATTGAAGCAATTCCTGCAGCCCGGGG | pAC23 cloning |
| AC483 | AGGGTCGACGGAAACCGATATCAAGCTTATCGATACCGTC | pAC23 cloning |
| AC484 | ATAAGCTTGATATCGGTTTCCGTCGACCCTCGG | pAC23 cloning |
| AC485 | CGGGCTGCAGGAATTGCTTCAATCCCACAAGGG | pAC23 cloning |
| AC499 | GTGGACCGGCGACCTCCCGGA | pAC28 cloning PaqCI cut site |
| AC500 | CTGCCTGCGCTTCAATCCCACAAGGG | pAC28 cloning PaqCI cut site |
| AC501 | TGCTTTGGTTTCAGACGAACCCTTGTGG | pAC29 cloning PaqCI cut site |
| AC502 | GGTGGGGAGGTCGCCGGTCCAC | pAC29 cloning PaqCI cut site |
| AC751 | CGAGGTCGACGGTATCGATACAGCAACTACTATGCGTTCGGAAGCCGAACTCTGCAGCCA | pAC53 cloning Insert *p.fdx* into HindIII cut site |
| AC752 | GTCGACGGAAACCGATATCATGGCTGCAGAGTTCGGCTTCCGAACGCATAGTAGTTGCTG | pAC53 cloning insert *p.fdx* into HindIII cut site |
| AC729 | GACTCACACCTGCGACTAAGCCACGGCCGAACTCCCCGACGAACTCCGGCCGGGAC | pAC70 cloning |
| AC730 | GTCACACACCTGCGTCAAAACGTCCCGGCCGGAGTTCGTCGGGGAGTTCGGCCGTG | pAC70 cloning |
| AC707 | GACTCACACCTGCGACTAAGCAAGACCGTCCCGTCGTGGCGGAGCGTCACCTCGTGGA | pAC59 cloning |
| AC708 | GTCACACACCTGCGTCAAAACTCCACGAGGTGACGCTCCGCCACGACGGGACGGTCTT | pAC59 cloning |
| AC413 | CGTCGCCGTATGTCTTCG | *mre11/rad50* spanning on genome |
| AC415 | CTCGTCAAGCAGTCGAGC | *mre11/rad50* spanning on genome |
| AC129 | GCGTTGTATTCGGGTATCTCGTAATC | spanning insertions on pTA232-based |
| AC130 | CGAGTACGCCTGCGACTACG | spanning insertions on pTA232-based |
| AC237 | NNNNNAAGGGTTCGTCTGAAACG | second PCR for acquisition (P1 and P2) |
| AC238 | NNNNNAAGGGTTCGTCTGAAACA | second PCR for acquisition (P1 and P2) |
| AC239 | NNNNNAAGGGTTCGTCTGAAACT | second PCR for acquisition (P1 and P2) |
| AC240 | NNNGTACAGCCGTCTACCC | second PCR for acquisition (P1 and P2) |
| AC241 | NNNNGTACAGCCGTCTACCC | second PCR for acquisition (P1 and P2) |
| AC242 | NNNNNGTACAGCCGTCTACCC | second PCR for acquisition (P1 and P2) |
| AC243 | NNNNNNGTACAGCCGTCTACCC | second PCR for acquisition (P1 and P2) |
| AC244 | CAAAGTGTTCCGGGAGGTCGCC | first PCR for acquisition (P1) |
| AC245 | CACGCCCTGCTGCCCGAA | first PCR for acquisition (P1) |
| AC247 | CAATATATTGCCTATGCCCGG | first PCR for acquisition (P2) |
| AC1089 | GAATTCCTCACCACATCGAG | first PCR for acquisition (P2) |
| AC683 | CCCGAATCAGGACGAAGAAC | PolD1 (HVO_0003) qPCR  Reference: (3) |
| AC684 | ATTTGAGGTGCTCGGAGAAC | PolD1 (HVO_0003) qPCR  Reference: (3) |
| AC1190 | TCGAAGACACAGCCGAGA | mre11 qPCR |
| AC1191 | CGAAATCACCCAGCGAACT | mre11 qPCR |
| AC1192 | GAAGACGTCGAGTCGCAAAT | rad50 qPCR |
| AC1193 | GCTCTCGTAGTTCGAAATCTCC | rad50 qPCR |
| AC1194 | TCACATCTTCGGCTGGAAAG | cas1 qPCR |
| AC1195 | CGTCGCTCCGTATTATCGTATG | cas1 qPCR |
| AC1196 | CCAAGAGGACTCGCATCTAC | cas2 qPCR |
| AC1197 | CTCACCGAAGAAGACCGAATAC | cas2 qPCR |
| AC1198 | TTGACCGTCGTCGTGAGATA | fen1 qPCR |
| AC1199 | TCTCCGAGGTGTCGTTTGA | fen1 qPCR |
| AC685 | TTGCGTACGCGGTATCTGT | HFPV-1 ORF2 qPCR  Reference: (3) |
| AC686 | AGCTTCTCCGCATCGTCTTT | HFPV-1 ORF2 qPCR Reference: (3) |
| AC444 | CGTTGCATCGGTTTCGGG | HFPV-1 fragment 1, sequencing PCR |
| AC445 | GGGTCAAGGTCATGCGCC | HFPV-1 fragment 1 |
| AC446 | GCCGCGGGGAAAGCCCGC | HFPV-1 fragment 2, sequencing PCR |
| AC447 | CGACGAACGAATACGGTGGTTCCTTC | HFPV-1 fragment 2, sequencing PCR |
| AC448 | GGCGAACGAATGGCGAAG | HFPV-1 fragment 3 |
| AC449 | CCCTTGCTCCCGAAGAAG | HFPV-1 fragment 3 |
| AC476 | GTCCGCGACGCGCTCTGCG | HFPV-1 pyrE2 |
| AC477 | GAAGCTCATCGCCGAAGC | HFPV-1 pyrE2 |
| AC478 | CAGCAACTACTATGCGTTCGGAAGCC | HFPV-1 pyrE2 |
| AC479 | TTAGCCGTCGGCGTCGGC | HFPV-1 pyrE2 |
| AC480 | TTATTTATAGAGTTCATCCATTCCG | HFPV-1 GFP |
| AC481 | CAGCAACTACTATGCGTTC | HFPV-1 GFP |
| AC608 | CGCAGTCCATCATCACGGACC | HFPV-1 ORF1^s^, ORF2^s^, ORF3^s^, ORF4^s^, ORF5^s^ |
| AC609 | GGAGGTCCGTGATGATGGACTG | HFPV-1 ORF1^s^, ORF2^s^, ORF3^s^, ORF4^s^, ORF5^s^ |
| AC610 | AATTCGGTGTCTCGTTAATCCTTTCAGGACCTC | HFPV-1 ORF1^s^ |
| AC611 | GAGGTCCTGAAAGGATTAACGAGACACCGAATT | HFPV-1 ORF1^s^ |
| AC620 | ATCATCACGAACGAGTAAAACACCGACCTATCG | HFPV-1 ORF2^s^ |
| AC621 | CGATAGGTCGGTGTTTTACTCGTTCGTGATGAT | HFPV-1 ORF2^s^ |
| AC622 | ATCACGATTTTATCCTAAGACCTGCTTGGTGGC | HFPV-1 ORF3^s^ |
| AC623 | GCCACCAAGCAGGTCTTAGGATAAAATCGTGAT | HFPV-1 ORF3^s^ |
| AC612 | GCGAACGACCAGAACTAAGACACCTATCTGCTC | HFPV-1 ORF4^s^ |
| AC613 | GAGCAGATAGGTGTCTTAGTTCTGGTCGTTCGC | HFPV-1 ORF4^s^ |
| AC624 | ACTCTCTCCCGAATCTAACTGAGCGAAGGAGAC | HFPV-1 ORF5^s^ |
| AC625 | GTCTCCTTCGCTCAGTTAGATTCGGGAGAGAGT | HFPV-1 ORF5^s^ |
| AC614 | GGCCGAGAGTCAGCAATCGG | HFPV-1 ORF6^s^, ORF7^s^, ORF8^s^, ORF9^s^ |
| AC615 | GGCCGATTGCTGACTCTCGG | HFPV-1 ORF6^s^, ORF7^s^, ORF8^s^, ORF9^s^ |
| AC616 | GACGAGGTGCTGGACTAAGTGATTGGAGAGTAC | HFPV-1 ORF6^s^ |
| AC617 | GTACTCTCCAATCACTTAGTCCAGCACCTCGTC | HFPV-1 ORF6^s^ |
| AC618 | CTGGACAGTCACGTCTAAGGGAAACAAATCGTG | HFPV-1 ORF7^s^ |
| AC619 | CACGATTTGTTTCCCTTAGACGTGACTGTCCAG | HFPV-1 ORF7^s^ |
| AC626 | ATCTTCGAAGACTATTAACACGAACGGCTCGCA | HFPV-1 ORF8^s^ |
| AC627 | TGCGAGCCGTTCGTGTTAATAGTCTTCGAAGAT | HFPV-1 ORF8^s^ |
| AC628 | GTCCTGAACCAACTCTAACGAATCCTAGAGCGA | HFPV-1 ORF9^s^ |
| AC629 | TCGCTCTAGGATTCGTTAGAGTTGGTTCAGGAC | HFPV-1 ORF9^s^ |
| AC630 | GAGGTCGTCCATGTGTAAGTTCCAGATGAACCG | HFPV-1 ORF10^s^ |
| AC631 | CGGTTCATCTGGAACTTACACATGGACGACCTC | HFPV-1 ORF10^s^ |
| AC632 | GGACGGAGCCAAGACCGGA | HFPV-1 ORF10^s^, ORF11^s^ |
| AC633 | GTGTTCCGGTCTTGGCTCCG | HFPV-1 ORF10^s^, ORF11^s^ |
| AC634 | AATTGGTCGGAGTGCTAAGGGATTGTCGTGTTG | HFPV-1 ORF11^s^ |
| AC635 | CAACACGACAATCCCTTAGCACTCCGACCAATT | HFPV-1 ORF11^s^ |
| AC425 | TGATGACGAATCCAACGAGCAG | HFPV-1 sequencing PCR |
| AC536 | GTCCGTTAGTCGGTAGACTCC | HFPV-1 sequencing PCR |

**Supplementary table 3.** Cloning and construction strategies for plasmids and HFPV-1 viruses constructed in this study.

| **Plasmid name** | **Cloning strategy** |
| --- | --- |
| pAC04 | PCR amplification of pTA232 using AC405/AC406, and of H119 gDNA using AC401/AC407 and AC408/AC409, followed by Gibson assembly of the three PCR fragments |
| pAC27 | PCR amplification of pTA232 using AC493/AC494, and of H119 gDNA using AC495/AC496 and AC497/AC498, followed by Gibson assembly of the three PCR fragments |
| pAC83 | PCR amplification of pTA232 using AC1069/AC1070, and of H119 gDNA using AC1071/AC1072 and AC1073/AC1074, followed by Gibson assembly of the three PCR fragments |
| pAC40 | PCR amplification of pTA232 using AC603/AC604, and annealed oligos AC601/AC602, followed by Gibson assembly of the two fragments |
| pMre11-Rad50 (pAC95) | PCR amplification of pTA232 using AC1135/AC1176, and of H119 gDNA using AC1138/AC1175, followed by Gibson assembly of the two PCR fragments |
| pFen1 (pAC42) | PCR amplification of pAC40 using AC589/AC592, and of H119 gDNA using AC590/AC591, followed by Gibson assembly of the two PCR fragments |
| pCas1-2 (pAC86) | PCR amplification of pAC40 using AC1083/AC1084, and of H119 gDNA using AC1085/AC1086, followed by Gibson assembly of the two PCR fragments |
| pAC23 | PCR amplification of H119 gDNA using AC452/AC486, followed by PCR amplification of that PCR product with AC484/AC485, and also PCR amplification of pTA232 of AC482/AC483, followed by Gibson assembly of the two PCR fragments |
| pAC28 | PCR amplification of pAC23 using AC499/AC500, followed by T4 PNK and T4 DNA Ligation (NEB, Cat #M0201 and #M0202) |
| pAC29 | PCR amplification of pAC23 using AC4501/AC502, followed by T4 PNK and T4 DNA Ligation (NEB, Cat #M0201 and #M0202) |
| pSpcNT (pAC53) | HindIII (NEB, Cat #R3104) treatment of pAC29, and annealed oligos AC751/AC752, followed by Gibson assembly of the two fragments |
| pSpc6 (pAC70) | PaqCI (NEB, Cat #R0745) treatment of pAC53, and annealed oligos AC729/AC730, followed by Gibson assembly of the two fragments |
| pSpc2971 (pAC59) | PaqCI (NEB, Cat #R0745) treatment of pAC53, and annealed oligos AC707/AC708, followed by Gibson assembly of the two fragments |
| **Virus name** | **Construction/cloning strategy** |
| HFPV-1 | Gibson assembly of three fragments ordered from FragmentGENE, Azenta Life Sciences, original sequences: (3) |
| HFPV-1 *(pyrE2)* | Gibson assembly of three fragments ordered from FragmentGENE, Azenta Life Sciences |
| HFPV-1* | Gibson assembly of four fragments ordered from FragmentGENE, Azenta Life Sciences |
| HFPV-1*-ORF1^s^ | PCR amplification of HFPV-1* using AC608/AC611 and also AC609/AC610, followed by Gibson assembly of the two PCR fragments |
| HFPV-1*-ORF2^s^ | PCR amplification of HFPV-1* using AC608/AC621 and also AC609/AC620, followed by Gibson assembly of the two PCR fragments |
| HFPV-1*-ORF3^s^ | PCR amplification of HFPV-1* using AC608/AC623 and also AC609/AC622, followed by Gibson assembly of the two PCR fragments |
| HFPV-1*-ORF4^s^ | PCR amplification of HFPV-1* using AC608/AC613 and also AC609/AC612, followed by Gibson assembly of the two PCR fragments |
| HFPV-1*-ORF5^s^ | PCR amplification of HFPV-1* using AC608/AC625 and also AC609/AC624, followed by Gibson assembly of the two PCR fragments |
| HFPV-1*-ORF6^s^ | PCR amplification of HFPV-1* using AC615/AC616 and also AC614/AC617, followed by Gibson assembly of the two PCR fragments |
| HFPV-1*-ORF7^s^ | PCR amplification of HFPV-1* using AC615/AC618 and also AC614/AC619, followed by Gibson assembly of the two PCR fragments |
| HFPV-1*-ORF8^s^ | PCR amplification of HFPV-1* using AC615/AC626 and also AC614/AC627, followed by Gibson assembly of the two PCR fragments |
| HFPV-1*-ORF9^s^ | PCR amplification of HFPV-1* using AC615/AC628 and also AC614/AC629, followed by Gibson assembly of the two PCR fragments |
| HFPV-1*-ORF11^s^ | PCR amplification of HFPV-1* using AC633/AC634 and also AC632/AC635, followed by Gibson assembly of the two PCR fragments |
